# MET inhibition exploits conserved regulatory dependencies in malignant rhabdoid tumors

**DOI:** 10.64898/2026.11.03.763704

**Authors:** Inés Sentís, Alex Cebria-Xart, Quirze Rovira, Elisa Melón-Ardanaz, Carlos J. Rodriguez-Hernandez, Samuel Morabito, Adrià Caballé, Irene Ruano, Andrea Urpi, Maria Caballero, Sara Garcia Palomino, Eva Rodriguez, Davide Maspero, Ginevra Caratù, Andrea Onieva, Domenica Marchese, Max Ruiz, Patricia Lorden, Israel Ramos, Cinzia E. Lavarino, Alicia Castaneda, Anna Pascual-Reguant, Jaume Mora, Juan C. Nieto, Holger Heyn, Alexandra Avgustinova

**Affiliations:** Institute for Research in Biomedicine (IRB Barcelona), The Barcelona Institute of Science and Technology, Barcelona, Spain; SJD Pediatric Cancer Center Barcelona (PCCB), Institut de Recerca Sant Joan de Déu (IRSJD), Esplugues de Llobregat, Barcelona, Spain; Centro Nacional de Análisis Genómico (CNAG), Barcelona, Spain; SJD Pediatric Cancer Center Barcelona (PCCB), Hospital Sant Joan de Déu, Esplugues de Llobregat, Barcelona, Spain; Universitat de Barcelona (UB), Barcelona, Spain; ICREA, Barcelona, Spain

**Author notes:** these authors contributed equally (Inés Sentís, Alex Cebria-Xart). these authors contributed equally (Quirze Rovira, Elisa Melón-Ardanaz). Co-corresponding authors: Alexandra Avgustinova, Holger Heyn.

**Keywords:** Tumor heterogeneity, Transcriptional plasticity, Single-cell multi-omics, Spatial transcriptomics, Gene regulatory networks, Biomarker-guided therapy, MET inhibition, Childhood cancer, Malignant rhabdoid tumor

## Abstract

Malignant rhabdoid tumors (MRTs) are lethal, yet genomically simple childhood cancers characterised by universal SWI/SNF dysregulation. Despite this shared molecular framework, the regulatory programs sustaining MRTs and how they wire actionable dependencies, remain poorly defined.

Here, we integrated single-cell multi-omic profiling and spatial transcriptomics across anatomically and molecularly diverse MRTs, generating a comprehensive multimodal MRT dataset at single-cell resolution. We leveraged this resource to identify candidate tumor-specific dependencies, which were subsequently functionally validated in vitro and in vivo in matched patient-derived models. We found that distinct malignant subpopulations are spatially patterned in complex environmentally shaped topologies, consistent with reciprocal interactions with stromal and immune compartments contributing to intratumoral transcriptional diversity. We identified recurrent transcriptional metaprograms conserved across patients and molecular subtypes, including a previously undescribed shared core rhabdoid program, as well as a metaprogram characterized by extracellular matrix interactions, and preferentially localized in tumor-stroma intermixed neighborhoods, supporting identification of niche conditioning cues. Mechanistically, we identified regulatory wiring driving these metaprograms, with the core shared rhabdoid programme coordinated by a circuit centred on TCF12, ZNF148 and CHD2.

Guided by target nominations from our integrated analyses, we performed a curated drug screen using patient-derived MRT models and validated HGF/MET signaling as a selective vulnerability in extrarenal rhabdoid tumors. Pharmacologic MET inhibition was cytotoxic in sensitive models and induced a complete response in vivo using a clinically approved compound. Importantly, MET expression in primary tumors correlated with response, supporting biomarker-guided patient stratification for immediate clinical translation.

Collectively, our findings define the molecular architecture of MRTs, reveal conserved regulatory programs beyond anatomical and molecular subtypes, and highlight a targetable signaling axis with immediate translational potential, establishing a framework for precision therapy in this lethal pediatric malignancy.

**Highlights:**

- Comprehensive multimodal atlas of malignant rhabdoid tumors (MRTs) integrating single-cell multi-omics, spatial transcriptomics, and in vivo functional validation in matched patient-derived models.
- Recurrent malignant transcriptional metaprograms are conserved across MRT subtypes and are spatially zonated within tumors, highlighting plasticity in response to environmental cues.
- A core rhabdoid regulatory circuit centred on TCF12/ZNF148/CHD2 coordinates the principal shared malignant program.
- Drug screening and functional validation identify core regulatory circuit-linked HGF/MET axis dependence in a subset of MRTs.
- Treatment with a clinically approved MET inhibitor induces striking complete response in vivo in MET-high tumors, supporting MET IHC-guided patient selection and immediate therapeutic translation.

## Introduction

Malignant rhabdoid tumors (MRTs) are highly deadly childhood cancers that most commonly affect infants and toddlers, with a median age at diagnosis below 2 years of age ^(1,2)^. Intensive multimodal therapy combining maximal surgical resection with chemotherapy and radiotherapy is the standard-of-care, achieving 5-year overall survival still below 30 % ^(3,4)^. Treatment-related toxicities amplified by the very young age underscore the need for effective and tumor-selective - yet currently completely lacking - targeted strategies.

A defining feature of rhabdoid tumors is their genomic simplicity: extreme clinical aggressiveness emerging from remarkably ‘quiet’ genomes. These cancers have an exceptionally low mutation rate, with biallelic functional loss of SMARCB1 (and very occasionally SMARCA4) acting as the key oncogenic driver event and, strikingly, the only recurrent mutation ^(5,6)^, rendering rhabdoid tumors reminiscent of a monogenic developmental disease. Mechanistically, SMARCB1 encodes a core subunit of the SWI/SNF chromatin remodeling complexes, and its loss drives widespread epigenetic dysregulation - disrupting enhancer control, lineage programmes, and cell-state regulation ^(1,7,8)^ - thereby enabling a single genetic insult to result in profound, genome-wide transcriptional rewiring. This biology is emblematic of childhood cancers more broadly, which generally harbor lower mutation burdens than adult tumors ^(9)^ and instead rely on developmental and epigenomic dysregulation for malignant transformation and maintenance of the oncogenic state ^(10,11)^.

Despite their similar genomic contexts, rhabdoid tumors are anatomically and molecularly heterogeneous. They occur across anatomical sites, including in central nervous system (atypical teratoid/rhabdoid tumors; ATRTs), kidney (rhabdoid tumors of the kidney; RTK), and extrarenal extracranial soft tissues (extrarenal rhabdoid tumors; ERRTs) ^(1,12)^. In ATRTs, comprehensive genome-wide DNA methylation and transcriptomic profiling have defined three major molecular subgroups - TYR, SHH, and MYC - with distinct transcriptional programmes and pathway activation ^(2,13,14)^. Outside the central nervous system, extracranial rhabdoid tumors (eMRT) likewise resolve into methylation-defined entities, including RTK-like and ERRT-like subtypes with distinct expression programmes ^(15)^. Notably, across intracranial and extracranial disease, molecular class and anatomical site are only partially coupled; related, but not interchangeable descriptors of rhabdoid tumor biology ^(12)^. Together, these observations suggest that a common initiating lesion can have divergent repercussions and govern distinct malignant regulatory programmes.

In parallel, heterogeneity extends to the non-malignant cell compartment. Despite their low tumor mutation burden, immune infiltration varies markedly across MRTs, with a subset - reported to be enriched among ATRT-MYC and extracranial cases - showing immune checkpoint–associated expression programmes ^(12,16)^. Clonally expanded intratumoral T cells support antigen-driven activation and raise the possibility that immune checkpoint inhibition (ICI) can be effective in selected patients ^(16,17)^.

Importantly, bulk profiling cannot determine how much of the apparent heterogeneity in MRTs reflects stable yet distinct malignant programmes versus spatially organized interactions with the local microenvironment. We therefore sought to resolve which malignant transcriptional states coexist within individual tumors, how these states are arranged within tissue topology, and whether spatial signaling and stromal context explain part of the observed cell state diversity, thereby enabling identification of core regulatory circuits and dependencies that transcend tumor anatomical site and subgroup label.

To address these questions, we performed multimodal single-cell and spatial profiling of a clinically and anatomically diverse MRT cohort, combining single-nucleus multi-omic profiling with spatial transcriptomics on matched sections and integrating these data with patient-derived models. This approach defines malignant cell states and infers their underlying regulatory programs, while mapping tumor– microenvironment (TME) interactions in situ. Leveraging cohort-level integration, we identify recurrent malignant transcriptional programmes shared across MRTs alongside context-skewed programmes that track with subtype and microenvironmental interactions, and we show that these programmes are spatially patterned into distinct tumor neighborhoods with characteristic signaling landscapes.

Finally, to translate our findings into actionable vulnerabilities we prioritized candidate pathways nominated by these integrated multi-omic analysis using a curated medium-throughput drug screen across patient-derived MRT cell models and controls, and then functionally validated prioritized candidate dependencies. This framework identified and functionally validated in vitro and in vivo a selective reliance on HGF/MET axis signaling in a subset of extracranial MRTs, with MET expression and activation predicting sensitivity. Together, our findings nominate MET inhibition as a biomarker-guided targeted treatment strategy for a defined subset of MRT patients.

## Results

### Multimodal profiling at single-cell resolution captures MRT diversity

To define both intra- and interpatient molecular and cellular diversity in a disease as genomically simple yet anatomically and molecularly heterogeneous as rhabdoid tumors, we performed multimodal characterization of a cohort consisting of 19 rhabdoid tumor patients treated at Sant Joan de Déu Children’s Hospital Barcelona between 2007 and 2024 (**Supplementary Table 1**). Samples included were ATRTs (n=5), RTKs (n=3), and ERRT (n=9). Accordingly, tumors originated from diverse anatomical locations, including brain (n=4), peripheral nerve (n=2), liver (n=1), kidney (n=3) and other soft tissues (n=7) (**Fig. 1a**), thus capturing the diversity in anatomical presentation inherent to this disease.

**Fig. 1.**
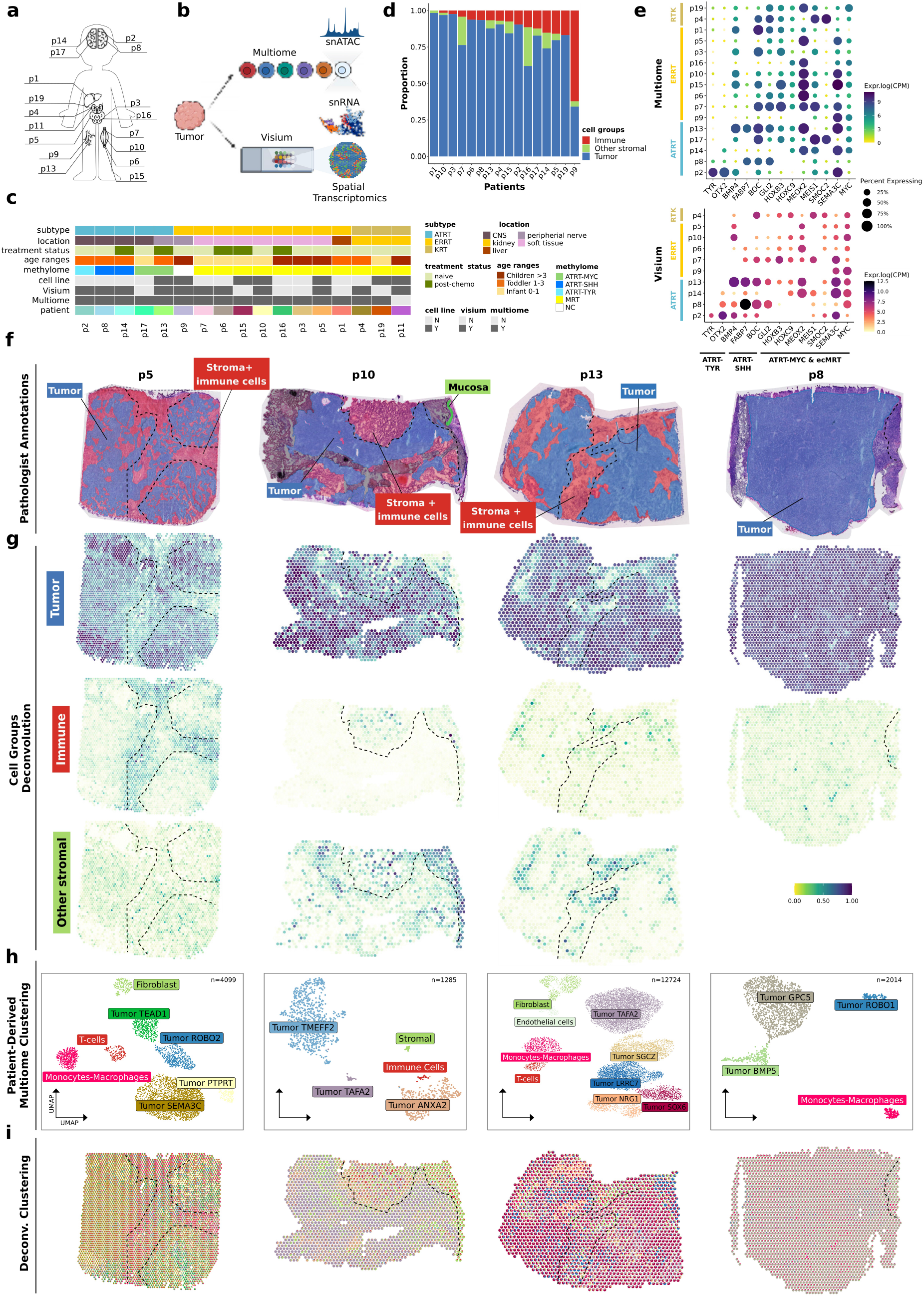
Multimodal profiling of MRT cohort highlights intra- and intertumor heterogeneity. **a**, Schematic illustration of anatomical location of patient samples. **b**, Schematic diagram of sequencing strategy. **c**, Summary table of cohort clinical information and data availability. **d**, Stacked barplot with the proportion of main cell groups (blue - tumor; red - immune; green - other stromal) per patient. **e**, Dot plot displaying the expression of canonical marker genes, with values representing log-transformed counts per million (logCPM) of pseudo-bulked data. Rows are categorized by patient clinical groups, while columns are organized by methylome-based molecular subgroups (following Chun et al., 2019). Top panel, estimated tumor fraction derived from the Multiome (snRNA assay only) dataset. Bottom panel, corresponding pseudo-bulk expression data derived from spatial Visium spots with high tumor purity (estimated deconvolution proportion > 0.5). **f**, Representative H&E-stained tissue sections from Visium slides of patients p5, p10, p13 and p8. Regions are delineated and color-coded based on expert pathologist’s annotation. **g**, Visium spots are color-coded based on the estimated proportions (ranging from 0 to 1, with low-value alpha transparency applied) of major cell groups derived from spatial deconvolution analysis. This visualization highlights the relative abundance and spatial distribution of distinct cell populations across the tissue architecture. Dashed lines delineate specific histological regions. **h**, UMAP visualization of patient-derived single-cell Multiome samples, with individual cells colored by their cluster annotations. **i**, Spatial mapping of patient-derived Multiome clusters onto Visium tissue sections. Each Visium spot is represented as a pie chart, where the slice sizes correspond to the deconvoluted proportions of the specific cell populations identified in panel **h**.

Frozen tumor specimens obtained from the Sant Joan de Déu Barcelona Children’s Hospital Biobank for Research (BHISJDI) were processed for single-nucleus multi-omic profiling (Chromium Single Cell Multiome ATAC + Gene Expression, 10X Genomics) and spatial transcriptomics (Visium, 10X Genomics) (**Fig. 1b**). To maximize similarity of the profiled specimens, serial histological sections were used whenever possible. From the 19 specimens processed, 17 passed our stringent QC criteria and were included in the cohort (**Supplementary Table 1, Fig. 1c**). We obtained high quality Multiome and Visium data from 16 and 11 patients, respectively. In addition, facilitating functional validation of our findings, cell lines were successfully established for 7 of the 19 primary patient samples.

Rhabdoid tumor identity was confirmed by a trained pathologist based on sample histological features including negativity for INI-1 staining (data not shown), and molecular subtypes were assigned by DNA methylation profiling (**Fig. 1c and Supplementary Table 1**). Genomic events leading to functional biallelic inactivation of SMARCB1 (and SMARCA4 in the case of p16) were confirmed for all patients using Sanger sequencing and Multiplex Ligation-dependent Probe Amplification (MLPA) (**Supplementary Table 1**). This anatomically and molecularly diverse cohort provides a representative framework to dissect malignant rhabdoid tumor heterogeneity.

### Spatially patterned malignant cell states emerge within heterogeneous rhabdoid tumor topologies

We first focused on the Multiome data. After removal of low-quality cells, doublets and ambient RNA contamination (see **Methods**), our Multiome dataset comprised 102,171 cells across 16 patients, with numbers ranging from 1,285 to 12,863 cells per sample (mean = 6,386, SD = 3,834; **Supplementary Table 1**). To characterize broad cellular composition, we initially resolved major cellular compartments on a per-patient basis (**Fig. 1d**). The analysis of both modalities revealed a clear separation of major cell lineages (tumor, immune and other stromal cells) within the first principal components for RNA and ATAC (**Supplementary Fig. 1a, Extended Data Fig. 1)**. The malignant cell fraction consistently expressed canonical subtype-specific rhabdoid marker genes ^(2,12–14)^ (**Fig. 1e; Supplementary Fig. 1a, Extended Data Fig. 1**).

To assess the spatial organization of these compartments, we analyzed matched tissue sections profiled with Visium. Visium spots with high tumor content also showed expression for canonical markers (**Fig. 1e**). Histopathological feature-based annotations by a trained pathologist delineated tumor cell-dominated areas from the infiltrating or adjacent stromal regions composed of fibrous connective tissue and immune cells, and served as a ground truth (**Fig. 1f**). The transcriptional profiles of the principal cell compartments were projected onto tissue space and showed high concordance with the expert annotation (**Fig. 1g, Extended Data Fig. 2a**). Inter-patient comparisons revealed substantial differences in tumor architecture and cellular heterogeneity between samples. Consistent with prior reports of variable immune infiltration ^(2,12,16)^, some tumors were highly infiltrated by immune cells (**Fig. 1g, left panel**), whereas others exhibited spatially segregated immune and stromal compartments, or were dominated by non-immune stromal cells (**Fig. 1g, middle panels**). A subset of tumors showed minimal infiltration by non-malignant cells, corresponding to a tumor-cell dominated, immune-cold phenotype (**Fig. 1g, right panel**). These observations highlight pronounced inter-patient heterogeneity in tumor tissue organization, in line with previously described diversity in immune landscapes of rhabdoid tumors ^(2,12,16)^. Notably, none of the sample’s descriptive characteristics accounted for the variations in stromal composition.

To further resolve intra-tumoral heterogeneity, we increased cellular resolution by sub-clustering each sample. This granular approach allowed for the distinction of myeloid, lymphoid, and stromal populations, while simultaneously characterizing a marked diversity in cellular states within the malignant cell compartment (**Fig. 1h, Supplementary Fig. 1a, Extended Data Fig. 1**). To confirm the rhabdoid identity of these tumor clusters, we quantified a pan-rhabdoid transcriptional signature (RT signature; see **Methods**). Malignant clusters exhibited systematically higher signature scores compared to the patient-matched non-malignant cell populations, validating their rhabdoid identity (**Supplementary Fig. 2a; Supplementary Table 2**). Such diversity in cellular states could in principle be driven by distinct mutationally-defined malignant cell clones, but is unexpected in broadly mutationally quiet rhabdoid tumors ^(6,9)^. Alternatively, spatially restricted niche signals, including extracellular matrix (ECM) composition and paracrine signaling interactions with non-malignant niche components, may contribute to the emergence of distinct malignant cell populations (**Supplementary Fig. 2b**). Consistent with spatially patterned microenvironmental cues, cell state deconvolution of the compendium of cellular states identified in our Multiome data within Visium spots using SPOTlight ^(18)^(see **Methods**) revealed pronounced spatial zonation across the tumor topographical space (**Fig. 1i, Extended Data Fig. 2b**). These observations suggest that local niche-specific signals generate distinct transcriptional states, resulting in spatially patterned malignant cell state diversity.

### Shared and context-dependent transcriptional metaprograms jointly shape rhabdoid tumor cell states

To identify recurrent transcriptional programs representing malignant cell states shared across rhabdoid tumors, we integrated the Multiome data with Harmony ^(19)^(see **Methods, Fig. 2a**), and unsupervised clustering yielded 14 clusters (**Fig. 2b**). Importantly, cells from different patients contributed to immune-related (clusters 3 and 11) and one non-immune stromal cluster (cluster 4), indicating effective correction of batch effects (**Fig. 2b**).

**Fig. 2.**
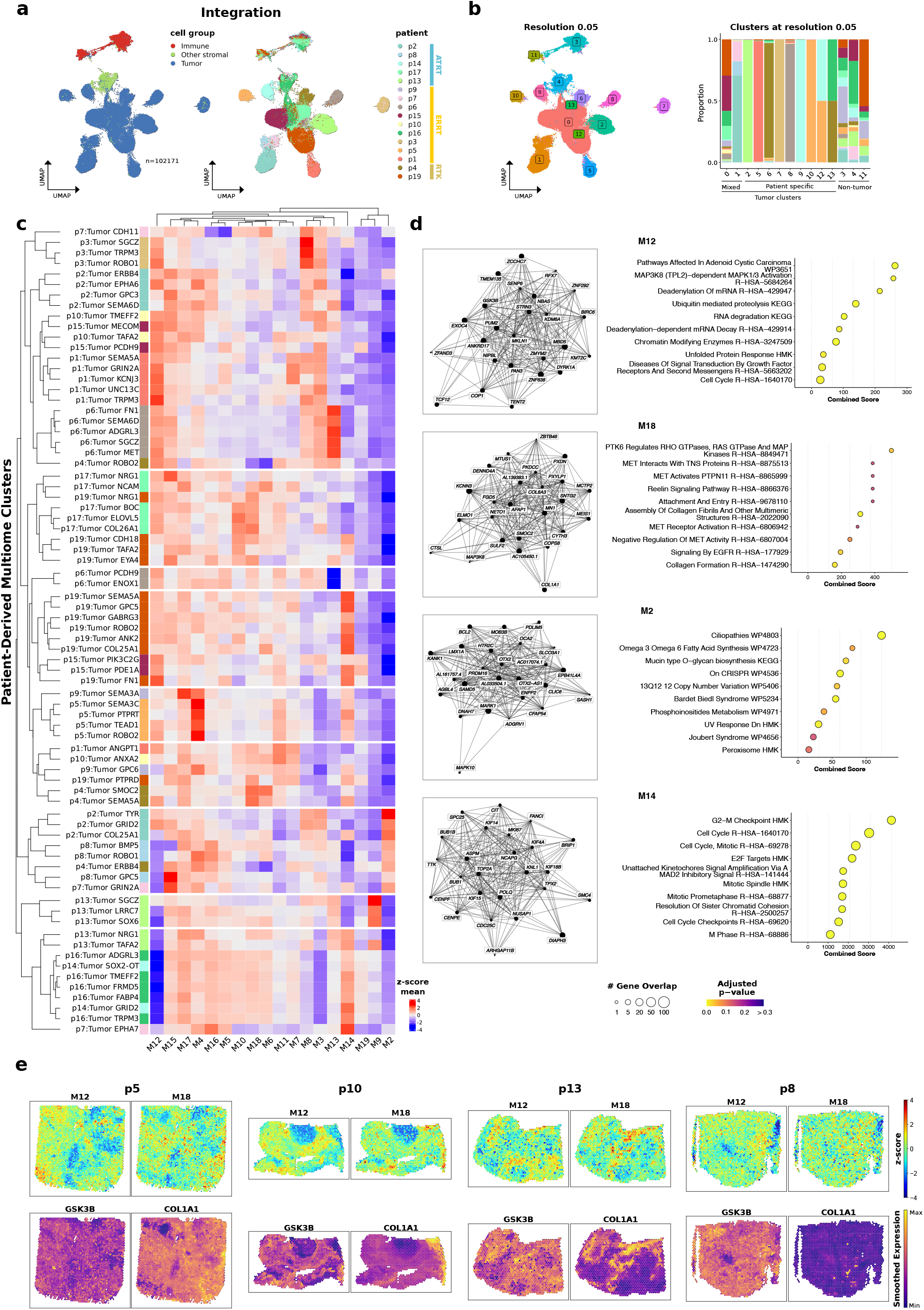
Integrated analysis of transcriptional metaprograms in MRTs. **a**, UMAPs of integrated rhabdoid patient samples. Left: colored by main cell groups; Right: colored by patient. **b**, Unsupervised clustering of integrated dataset. Left, UMAP from **a** colored by clusters obtained at 0.05 resolution. Right, stacked bar plot depicting the distribution of patient-specific cell proportions within each cluster at matched resolution. **c**, Heatmap summarizing the average activity of co-expression modules within tumor populations from patient-derived Multiome clusters (see also Supplementary Fig.1). Values are z-scored row-wise to highlight relative expression differences, with rows organized by hierarchical clustering. **d**, Left, gene network connectivity plots for metaprograms M12, M18, M2, and M14. Right, corresponding ORA dot plots for each module. Y-axis displays the top enriched biological terms (HMK:Hallmarks); x-axis indicates the combined score. Dot size represents the gene count overlap, and color indicates the adjusted p-value. **e**, Top, spatial distribution of M12 and M18 activity, calculated as metaprogram signatures and z-score per sample. Bottom, smoothed spatial expression of hub genes *GSK3B* (M12) and *COL1A1* (M18); values were imputed to enhance visualization of spatial patterns and regional colocalization.

In contrast, most malignant cell clusters were largely patient-specific, with the apparent idiosyncratic biology of malignant cells increasing with clustering resolution (**Supplementary Fig. 3a**). Consistently, principal component (PC) analysis of the integrated embedding revealed that PC1 captured biological variation, whereas PC2-5 were indeed driven by patient identity (**Supplementary Fig. 3b**). Nevertheless, clustering at lower resolution (0.05) highlighted broadly similar transcriptional programs shared by multiple patients. Cluster 0 showed contribution from 14/16 patients and was enriched for pathways related with oxidative phosphorylation and elevated metabolic activity (**Supplementary Fig. 3c**). This is consistent with previous reports of reliance of RT on this pathway ^(20)^. Additionally, spatial analysis confirmed that the oxidative phosphorylation program sharply delineated tumor-dominated regions, which exhibited high activity, from adjacent stromal regions (**Supplementary Fig. 3d; Extended Data Fig. 3a**). This pattern was conserved across all samples in our cohort, independent of anatomical location or molecular subtype, highlighting a robust metabolic distinction between malignant and non-malignant compartments and highlighting a potential specific yet subtype-agnostic vulnerability of rhabdoid tumors.

Cluster 1, on the other hand, was predominantly constituted by cells derived from cranial ATRT-SHH and ATRT-TYR tumors and displayed transcriptional profiles consistent with previously reported ATRT programs, including ciliopathy-related genes and *OTX2* and *TYR* expression ^(14,21,22)^(**Supplementary Fig. 3c**). This cluster highlights relatively closer transcriptional similarity between ATRT-TYR and ATRT-SHH tumors, and divergence from ATRT-MYC and eMRTs in agreement with bulk methylation and transcriptomic analyses ^(12)^.

To further dissect inter-patient transcriptional heterogeneity, we inspected the variability within the integrated tumor fraction across patients. Principal component analysis (PCA) showed that the first principal component (PC1) separated ‘ATRT-like’ versus ATRT-MYC/eMRTs (**Supplementary Fig. 3e**). Examination of PC1 gene loadings revealed that this variance was driven by the expression of specific markers. Intracranial ‘ATRT-like’ samples were defined by genes such as *TYR* and *CLIC6*, whereas the ATRT-MYC/eMRT axis was characterized by the mesodermal transcription factor *MEOX2* (**Supplementary Fig. 3f**). Inspecting the expression of highly variable genes (HVGs), we found growth factor-mediated signaling pathways, including multiple receptor tyrosine kinase pathways, such as EGF/EGFR, FGF/FGFR and PDGF/PDGFR signaling, consistent with prior reports in MRTs ^(23–25)^ (**Supplementary Fig. 3g**). In addition, hepatocyte growth factor (HGF) emerged as one of the top PC1 loadings, together with its cognate receptor, MET, highlighting the HGF/MET pathways as a novel potential vulnerability of rhabdoid tumors. Further biological categories of HVGs included ECM and cell adhesion, as well as transcription factors (TFs), suggesting that microenvironmental cues and intrinsic regulatory networks jointly shape cellular heterogeneity in rhabdoid tumors. Finally, HVGs also corresponded to potential immunotherapy targets, including oncofetal antigens ROBO1 ^(26,27)^ and GPC3 ^(28–30)^ as well as other glypicans (GPC5, GPC6), and LGR5 ^(31)^, the latter a well-established marker of stem cell populations in normal and malignant tissues ^(32)^. Together, these observations highlight potential targets for future therapy development in rhabdoid tumors.

While PCA highlighted the major axes of transcriptional divergence between broader groups of rhabdoid tumors, we next set out to identify transcriptional programs conserved across rhabdoid tumor subtypes. To systematically identify co-expressed gene networks that capture shared rhabdoid biology, we applied hdWGCNA in the tumor fraction ^(33)^(**Fig. 2c,d, Supplementary Fig. 3h**). We identified 19 co-expressed gene modules, herein designated as transcriptional metaprograms (M), many of which were subtype agnostic and shared across multiple patients (**Supplementary Table 3, Supplementary Fig. 3h**).

M2 was predominantly enriched in ATRT-SHH/ATRT-TYR-like tumors and pathway analysis revealed enrichment for ciliopathy-related programs (**Fig. 2d**), as previously described in ATRTs (14,21,22). In addition, the top hub genes of M2 included key drivers of ATRT biology, such as *OTX2, BCL2* and *CLIC6*, further supporting M2 as an ATRT-like transcriptional metaprogram.

To identify a core shared MRT transcriptional program, we focused on metaprograms that were broadly expressed across patients. In that regard, M12 represented a conserved transcriptional program highly active within the majority of malignant subpopulations (**Fig. 2c; Supplementary Fig. 3h, Extended Data Fig. 4**). Over representation analysis (ORA) identified pathways with clear significance for rhabdoid tumor biology. For instance, enrichment for ‘Chromatin modifying enzymes’ likely reflects compensation for functional loss of SMARCB1, which leads to severe chromatin dysregulation ^(1)^. In addition, ‘ubiquitin-mediated proteolysis’ has recently been recognized to be important for the clearance of incorrectly assembled SWI/SNF complexes in the context of SMARCB1 loss ^(34)^. High levels of ‘Unfolded protein response’ have also been described in rhabdoid tumors ^(35)^. Notably, one of the hub genes with the highest connectivity within M12 was *GSK3B*, a key regulator of WNT developmental signaling. While GSK3B phosphorylation, rather than expression, is more informative with respect to WNT pathway activation, we note the possibility that WNT activity may be involved in the regulation of the core rhabdoid transcriptional metaprogram.

Notably, M12 underpinned biologically meaningful separation in unsupervised hierarchical clustering of patient-derived tumor clusters by metaprogram composition, dividing tumor cell populations into M12-high and M12-low clusters (**Fig. 2c**). The M12-low cluster further segregated into a subcluster defined by high M2, composed of subpopulations from ATRTs (p2 and p8), as well as extracranial tumors (p4 and p7). Additionally, M14 drove the convergence of tumor cell subpopulations from different patients within both M12-high and M12-low clusters. M14 represented a proliferation-associated transcriptional metaprogram, enriched for cell-cycle-related pathways and hub genes including *MKI67* and *CDC25C*. M14 expression at patient-level correlated with an external proliferation signature across tumors ^(36)^, and Ki67 immunohistochemistry further confirmed these findings at the protein level (**Fig. 2d, Supplementary Fig. 3i,j**).

Transcriptional programs shared across patients further identified M18, enriched for MET receptor signaling, as well as ECM deposition and cell adhesion-related pathways (**Fig. 2c,d**). Hypothesizing M18 to be preferentially enriched at tumor-stroma interfaces, we mapped M12 (core rhabdoid program), M18 (ECM-related program), and representative hub genes (*GSK3B* and *COL1A1* for M12 and M18, respectively) in space. Indeed, M12 and M18 displayed distinct spatial localization, with M18 localizing to regions associated with higher non-immune stromal content, consistent with the expected TME crosstalk (**Fig. 2e; Extended Data Fig. 3b**).

To determine whether the transcriptional metaprograms identified in our single-cell multi-omic dataset reflected recurrent features of rhabdoid tumor biology, we evaluated their activity in an independent bulk RNA-sequencing cohort from the St. Jude Cloud pediatric cancer resource ^(37)^. We computed our gene metaprograms as signature scores across the compiled 3,029-sample pediatric pan-cancer dataset, and compared ATRTs and eMRTs with other major pediatric malignancies.

Consistent with our single-cell analyses, M2 showed marked specificity for ATRTs, with substantially lower activity in other pediatric tumor types (**Supplementary Fig. 3k**). In contrast, M12 was elevated in both ATRT and eMRT samples compared to the wider pediatric cancer cohort. Within this rhabdoid-specific axis, M12 activity was slightly more pronounced in eMRTs than in ATRT, directly recapitulating the pattern observed in our single-cell multi-omic dataset.

Together, our single-cell analyses reveal a recurrent transcriptional architecture in rhabdoid tumors, integrating a shared core rhabdoid program (M12), subtype-associated programs (M2), stromal-associated states (M18), and proliferative activity (M14). Analysis of an independent pediatric pan-cancer bulk RNA-sequencing cohort validated the ATRT specificity of M2 and the enrichment of M12 across rhabdoid tumors, indicating that these metaprograms capture conserved biology across cohorts. Thus, single-cell metaprogram analysis resolves the apparent heterogeneity of MRTs into biologically interpretable axes that reflect core identity, subtype specification, proliferation and microenvironmental context.

### Gene regulatory circuits at the basis of transcriptional metaprograms

To determine whether the identified co-expression metaprograms were underpinned by coordinated gene regulatory networks (GRNs), we used SCENIC+, a multi-omic framework that integrates transcriptomic and chromatin accessibility data to infer eRegulons (comprising TFs, cis-regulatory elements, and associated target genes)^(38)^.

At cohort level, we found genes in rhabdoid metaprograms to be co-regulated by defined sets of TFs (**Fig. 3a; Supplementary Fig. 4a,b**). Specifically, the M12 metaprogram was driven by a regulatory circuit involving TCF12, ZNF148, and CHD2 (**Fig. 3b; Supplementary Table 4**). Notably, TCF12 itself ranked among the top hub genes of the M12 metaprogram (**Fig. 2d**), directly linking metaprogram structure with its inferred regulatory circuitry. All three eRegulons were enriched in the malignant rhabdoid compartment, with CHD2 showing the strongest specificity for the malignant cell fraction (**Supplementary Fig. 4c**). Further, the M14 metaprogram was governed by TFs essential for cell cycle regulation (e.g. E2F7, E2F8), which directly correlated with the G2/M and S-phase states identified in the malignant populations (**Fig. 3a,c; Supplementary Fig. 4d,e**). In contrast, the M2 metaprogram was driven by eRegulons centered on TFs previously established as core drivers of ATRT biology such as OTX2 ^(14)^ (**Fig. 3a,d,e**). Importantly, eRegulon and metaprogram activities show strong agreement at the Multiome cluster level, consistent with metaprograms reflecting underlying GRN wiring. Moreover, we observed that eRegulons associated with the same metaprogram exhibited highly correlated activity profiles, indicating that these TFs are jointly active within the same cellular states. Despite shared activation patterns, downstream target genes for each eRegulon were highly divergent, overall suggesting a coordinated but non-redundant regulatory circuit (**Supplementary Fig. 4f,g**).

**Fig. 3.**
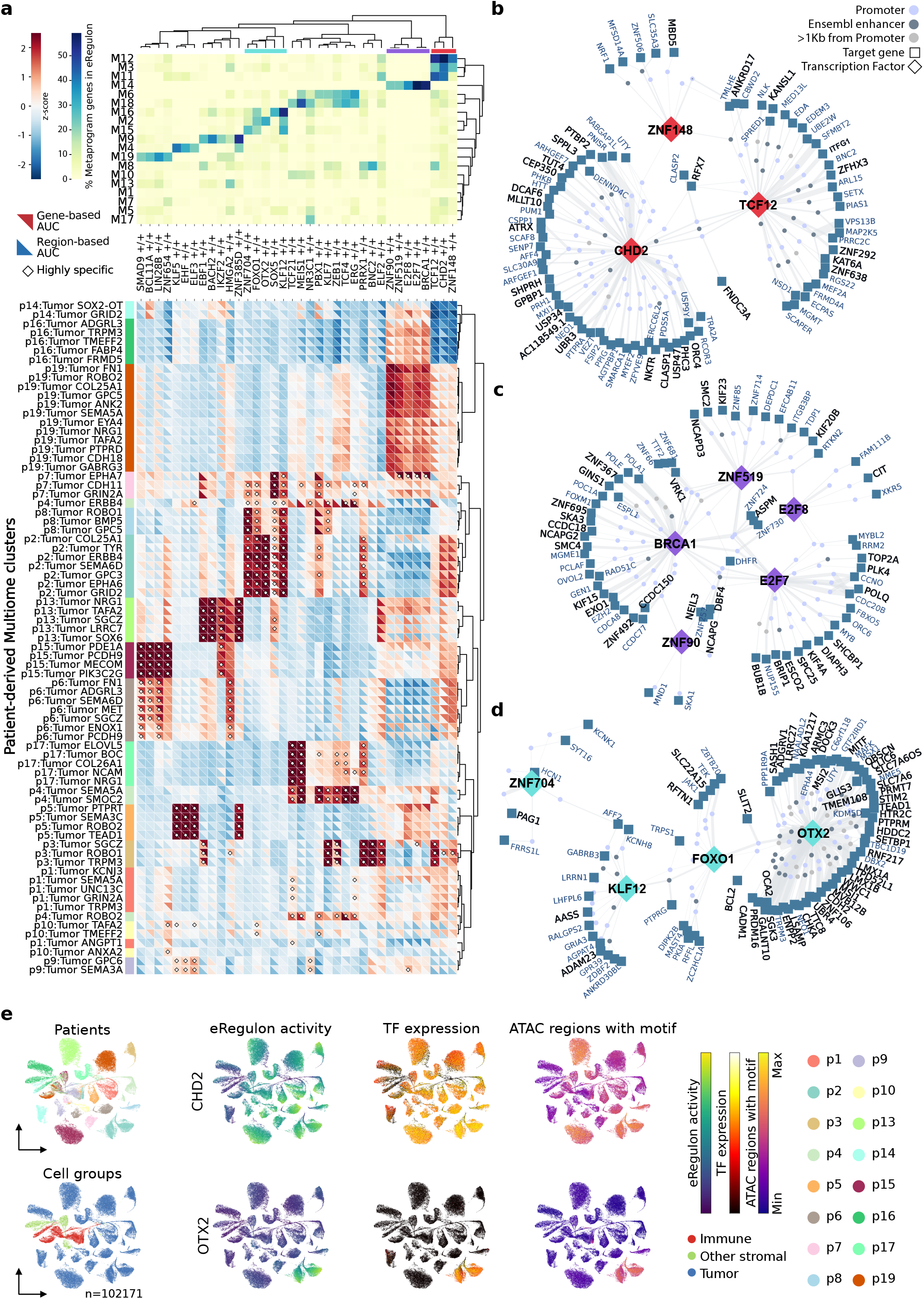
Gene regulatory circuits underlie transcriptional metaprograms. **a**, eRegulons identified by SCENIC+ and overlapping with metaprogam genes. Top: percentage of metaprogam genes that overlap with eRegulon target genes. Bottom: gene (upper-triangle) and region (bottom-triangle) eRegulon score for each patient-derived Multiome cluster. Clusters in which an eRegulon is considered highly specific based on regulon-specificity score are marked with a diamond. **b-d** Gene regulatory networks of selected eRegulons highlighted in **a** as overlapping with specific metaprograms (b-M12, c-M14, d-M2). Genes overlapping between indicated eRegulon and metaprograms are highlighted in black. Dots representing linking regions are classified as *Promoter* if they are within 1Kb of the TSS, *Ensembl enhancers* if they overlap with enhancers in Ensembl regulatory feature annotations, or *>1Kb from Promoter* for all others. Connecting line width represents the importance of the TF as a predictor of gene expression. **e**, UMAP dimensionality reduction based on gene-based AUC eRegulon scores. Left, colored by patient and main cell groups. eRegulon activity, TF expression and ATAC signal of linked peaks are shown for representative eRegulons CHD2 and OTX2.

Together, integration of chromatin accessibility with single-cell transcriptional states resolved metaprogram-associated TF networks, revealing that rhabdoid metaprograms are anchored in coherent yet modular and non-redundant gene regulatory circuitry. This regulatory framework enables prioritization of candidate dependencies and downstream effector programs, ultimately uncovering potential therapeutic vulnerabilities embedded within recurrent rhabdoid tumor states.

### Spatial niches delineate recurrent cell–cell communication in rhabdoid tumors

Given that several of the identified transcriptional metaprograms, most prominently M18, reflected cell-cell communication, we next used the spatial resolution of our cohort to systematically map recurrent ligand-receptor interactions within the rhabdoid tumor compartment, as well as between malignant cells and the adjacent stroma. Therefore, we deconvoluted spatial transcriptomics spots into the main cell compartments (malignant, immune, other stromal; see **Fig. 1; Extended Data Fig. 2b**) and analyzed spatial neighborhoods, defined as spots within a 100 µm radius. To quantify local cellular diversity, we calculated Shannon Entropy across Visium spots: high entropy regions corresponded to mixed tumor-stromal neighborhoods, whereas low-entropy regions marked homogeneous tumor or stromal areas (**Fig. 4a**). This analysis revealed three classes of tumors - those with low, medium, or high compositional complexity - and identified five recurrent spatial architectures hereafter referred to as ‘niches’ (**Fig. 4b,c**). Tumor-dominated (N2) and tumor high (N0) niches were almost entirely composed of malignant cells, with N0 containing a smaller contribution of non-malignant cells. Mixed neighborhoods (N1) contained a larger proportion of immune cells, while stromal-enriched (N3) and immune-enriched (N4) neighborhoods were dominated by the respective non-malignant compartments. These niches provided a spatial framework to relate spatial heterogeneity to recurrent cell-cell interactions and signaling programs across rhabdoid tumors.

**Fig. 4.**
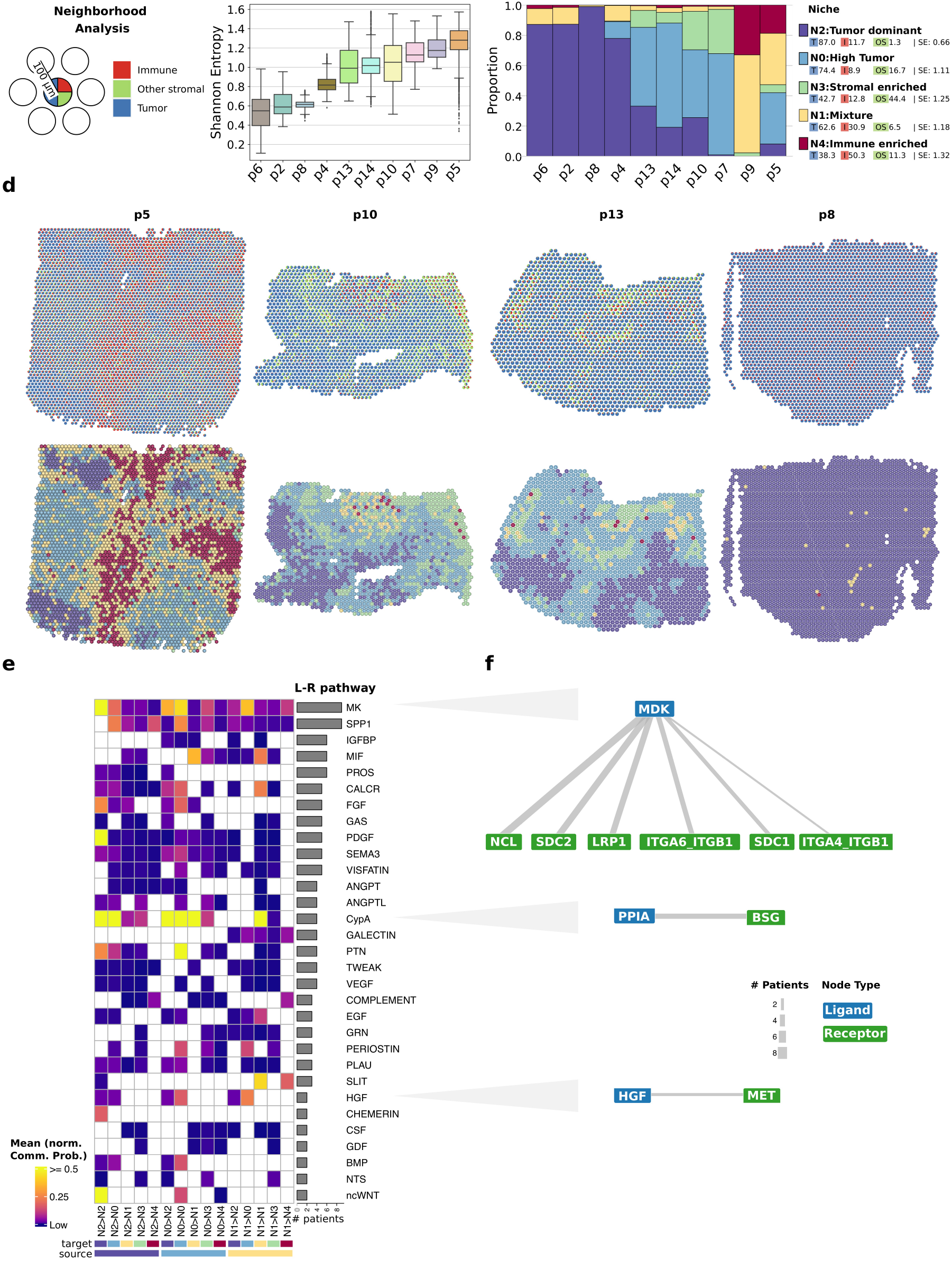
Ligand-Receptor signaling in spatial niches. **a**, Schematic of the neighborhood analysis approach using cell group proportions per spot and evaluating heterogeneity in a radius of 100 µm. **b**, Box plots displaying the distribution of Shannon Entropy scores per spot, stratified by patient and ordered by median values. **c**, Stacked bar plots illustrating proportions of spatial spots assigned to specific spatial niches across the patient cohort. **d**, Top, spatial deconvolution maps where each spot is represented as a pie chart of cell group proportions. Bottom, corresponding spatial maps with spots color-coded by their assigned histological niche. **e**, Heatmap illustrates the average communication probability between specific source and target histological niches across the patient cohort (source niches with high tumoral content only). For each patient, raw signaling probabilities (derived from CellChat) were normalized to the maximum observed value to allow for inter-sample comparison. The heatmap displays the mean of these normalized probabilities for each ligand-receptor pathway across all patients. To emphasize biological signals, the color scale is capped at 0.5 (maximum intensity); white color represents no significant communication drawn by CellChat. Right annotation bar plot indicates the total number of patients in whom significant communication was detected for that specific pathway. **f**, Diagram representing ligand and partner receptor(s) for selected pathways (MK, CypA, HGF). Edge size represents the number of patients where the indicated signaling interaction was inferred.

Spatial visualization of these niches confirmed the heterogeneity in cell composition between tumors and revealed clear zonation and topological organization. As predicted, M18, which we previously identified as an edge-localized metaprogram with strong stromal signaling (**Fig. 2e**), was more strongly detected in stomal-enriched neighborhoods relative to tumor-dominated regions, supporting its role in TME interactions (**Fig. 4d**). We next performed cell-cell communication analysis using CellChat (see **Methods**) to identify recurrent signaling axes across patients. We initially focused on interactions initiated by malignant cells (ligands expressed in N0, N1 and N2 niches; **Fig. 4e**). This revealed multiple recurrent interactions, most prominently midkine (MDK), MIF, CypA, SPP1 and HGF (**Fig. 4e,f**; **Supplementary Fig. 5a,b**). Notably MIF, SPP1 and MDK were also detected as ligands secreted from immune- and stromal-dominated neighborhoods, indicating that these signals operate in multiple spatial contexts. In addition, HGF was secreted by tumor cells, yet HGF/MET signaling was strongest in mixed neighborhoods, suggesting that both autocrine and paracrine signalling may contribute to tumor growth (**Supplementary Fig. 5a,b**).

We next focused on characterizing the stromal landscape in more detail. The harmonized immune compartment (**Supplementary Fig. 5c**) was dominated by M2-like macrophages, with variable T cell infiltration across tumors (**Supplementary Fig. 5d,e**)^(2,12,16)^. The non-immune TME contained fibroblasts expressing high levels of *SULF1*, consistent with TGFβ signaling in stromal-enriched niches (39,40) and endothelial and mural cells, related to the high levels of angiogenesis in rhabdoid tumors ^(41)^(**Supplementary Fig. 5f-h)**. Altogether, the TME of rhabdoid tumors is characterized by broadly anti-inflammatory features and a pro-tumorigenic milieu.

Together, these findings link stromal-associated metaprograms to recurrent spatial niches and signaling axes, indicating that part of the apparent transcriptional heterogeneity of rhabdoid tumors reflects structured interactions between malignant cells and a pro-tumorigenic microenvironment.

### Curated drug screen identifies HGF/MET axis as a selective vulnerability in ERRTs

Finally, to translate the regulatory and signaling features identified by our multi-omic analyses into actionable vulnerabilities, we designed a curated medium-throughput drug screen focused on pathways nominated by our data. We profiled our newly established patient-derived rhabdoid tumor cell models (n = 7), together with commercial ATRT cell lines (n=2; CHLA-02 and CHLA-05), and non-rhabdoid control cells - mesenchymal stromal cells (HFF-1), normal epithelial cells (HEK293) and neural-like cells (SH-SY5Y) - to assess tumor specificity. The screen included 44 compounds targeting candidate dependencies linked to activated developmental programs, TME-associated signaling axes, as well as previously reported rhabdoid tumor vulnerabilities. More specifically, we targeted (1) the OXPHOS pathway (see **Supplementary Fig. 3c,d, Extended Data Fig. 3a**); (2) developmental signaling axes, including WNT signaling (GSK3B core of M12; see **Fig. 2**) and BMP signaling (see **Fig. 1e, Supplementary Fig. 3f**); and (3) cell-cell interactions, including HGF/MET and MDK (see **Fig. 4, Supplementary Fig. 5a,b**). To evaluate relative specificity, we also included six conventional chemotherapy agents, including doxorubicin, etoposide and vincristine, which form part of standard-of-care treatment for patients with rhabdoid tumors in the EU-RHAB protocol.

For the drug screen, growth inhibition after 72h was assessed at two concentrations: 100 nM, selected to approximate a clinically relevant exposure (**Fig. 5a**), and 1 μM, to capture pathways that may become targetable with optimized inhibitors (**Supplementary Fig. 6a**). As expected, classical chemotherapy agents showed strong activity at low concentrations but limited selectivity for rhabdoid cells as compared to control cells, consistent with their known systemic toxicity profile in patients ^(3)^. Similarly, OXPHOS inhibitors broadly decreased viability among all tested cell models, including controls, at both concentrations. In contrast, the epigenetic agents decitabine and panobinostat displayed moderate efficacy and selectivity for rhabdoid cells; both compounds have previously been trialed in rhabdoid tumor patients, albeit with limited clinical benefit ^(42)^(NCT04897880). Notably, selectivity decreased for most agents at the higher concentration, underscoring the importance of dose to preserve tumor selectivity and avoid broad, non-specific cytotoxicity.

**Fig. 5.**
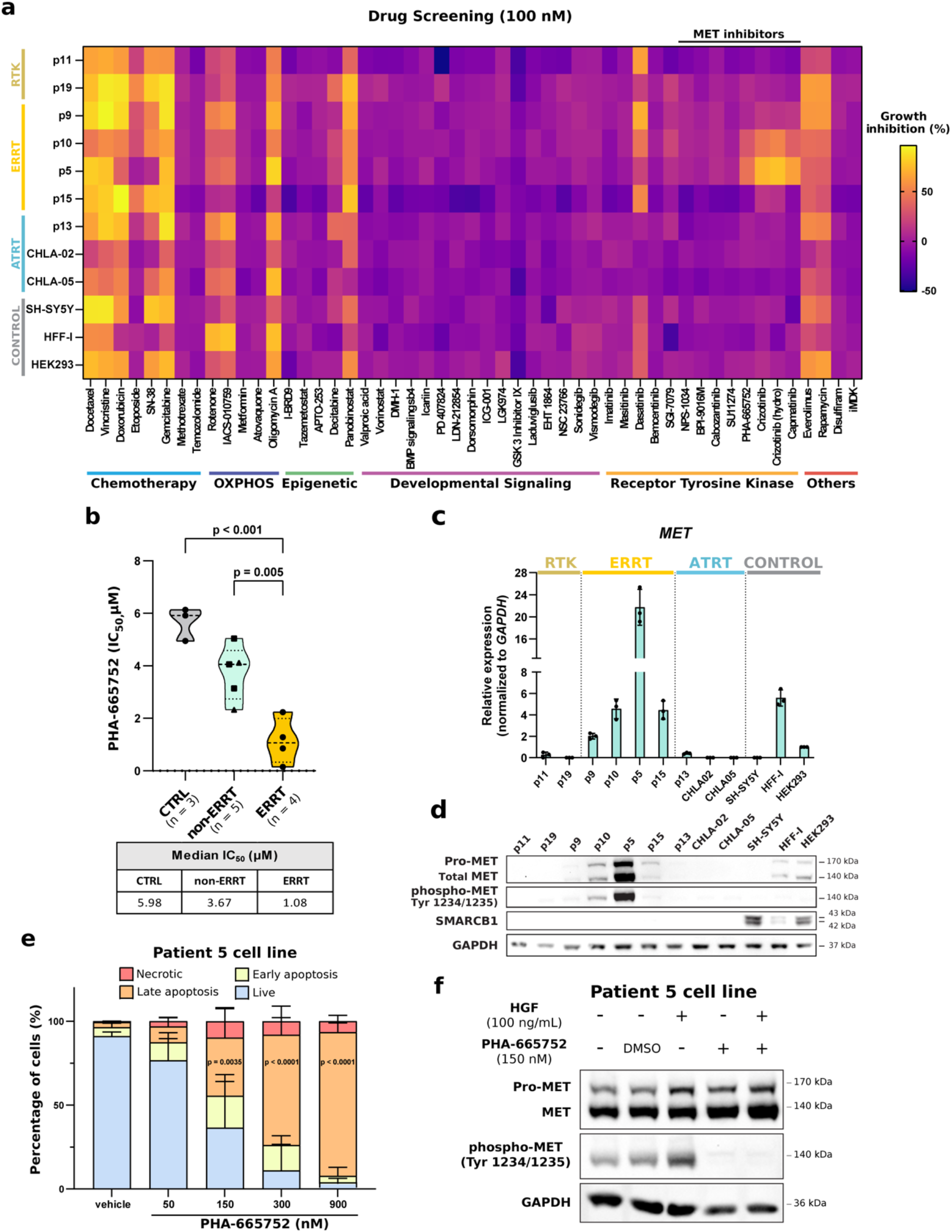
HGF/MET axis as a vulnerability in a subset of MRTs. **a**, Heatmap depicting the percentage of growth inhibition across a panel of patient-derived (RTK, ERRT, ATRT) and control (SH-SY5Y, HFF-1 and HEK293) cell lines. Straight line indicates inhibitors targeting MET. Data represents the mean of three technical replicates following 72 hours of treatment at 100 nM. **b**, Violin plot showing the distribution of *LogIC*_*50*_ values for the MET inhibitor PHA-665752 across experimental groups. Statistical significance was determined by one-way ANOVA followed by Tukey’s multiple comparisons test (n = 3 independent experimental replicates). Calculated median *IC*_*50*_ values are shown in the table (below). **c**, Relative mRNA expression of *MET* in tumor cell lines assessed by RT-qPCR (n = 3 independent experimental replicates). **d**, Whole protein lysates from patient-derived rhabdoid tumor models and control cell lines probed for the indicated proteins. GAPDH was used as a loading control. **e**, Quantification of live, necrotic, early and late apoptotic patient 5 cells after treatment with vehicle or PHA-665752 at the indicated concentrations for 24 hours. Statistical significance was determined by two-way ANOVA followed by Dunnett’s multiple comparisons test (n = 3 independent experimental replicates). **f**, Western blot for the indicated proteins in patient 5 cells supplemented or not with 100 ng/mL human HGF recombinant protein and/or 150 nM PHA-665752 for 3 hours. GAPDH was used as a loading control.

In contrast, targeting specific developmental signaling pathways produced negligible effects in this assay. Compounds modulating canonical or non-canonical WNT signaling showed no anti-proliferative effects in any of the tested cell models. Although the short assay timeframe may not capture slower phenotypes such as differentiation or cell state remodeling, the absence of acute growth inhibition suggests that these axes are unlikely to represent primary dependencies in rhabdoid cells. Similarly, MDK inhibition did not produce detectable effects under monoculture conditions, suggesting that this spatially nominated ligand-receptor axis may require multicellular context to reveal functional consequences.

Interestingly, inhibition of BMP receptor signaling using the ALK2 inhibitor LDN-212854 showed high specificity for ATRTs and neural-like SH-SY5Y cells at high concentrations (**Fig. 5a, Supplementary Fig. 6a**). Clinically actionable ALK2 inhibitors are not yet available, suggesting that this pathway could represent a target for future therapeutic development.

The most striking and specific vulnerability identified by our drug-screen was MET inhibition. The ERRT subset of rhabdoid models was sensitive to MET blockade across mechanistically distinct inhibitors (including capmatinib, crizotinib, and PHA-665752), consistent with tumor cell reliance on HGF/MET pathway activity. These compounds induced near-complete growth inhibition (> 80 %) in ERRT cells within the tested timeframe, while ATRT models and non-rhabdoid controls were largely unaffected. This selective sensitivity, together with the clinical availability of multiple FDA-approved MET inhibitors, nominates MET as an actionable target with potential for near-term clinical translation.

To validate these findings, we tested PHA-665752 across a broader concentration range. ERRT cell lines had significantly lower IC_50_ values (median = 1.08 μM; 95 % Confidence Interval (CI) 0.15 – 2.3 μM) than control and non-ERRT models (median = 5.98 μM and 3.67 μM, with CI 5.02 – 6.16 μM and 2.33 – 5.1 μM, respectively)(**Fig. 5b, Supplementary Fig. 6b**), supporting greater dependence on HGF/MET signaling. Accordingly, ERRT-derived models exhibited higher pro-MET/total MET and elevated phospho-MET as compared to non-ERRT counterparts (**Fig. 5c, d; Supplementary Fig 6c**). Indeed, within ERRTs, higher MET expression strongly predicted sensitivity to MET inhibition (**Supplementary Fig. 6d**). Mechanistically, Annexin V staining showed dose-dependent apoptosis in ERRT, but not ATRT or control cells, indicating ERRT-specific cytotoxicity of MET inhibition (**Fig. 5e, Supplementary Fig. 6e**). Consistent with pathway dependence, in ERRT cells MET phosphorylation was further increased by recombinant HGF and completely abrogated by pharmacological MET inhibition without altering total MET level (**Fig. 5f**). To test ligand dependence, we neutralized HGF with ficlatuzumab, a clinical-stage humanized anti-HGF antibody. ERRT cell lines showed a dose-dependent loss of viability, whereas RTK, ATRT and control lines were largely unaffected by HGF neutralization (**Supplementary Fig. 6f**). Together with the small molecule data, these results support HGF/MET autocrine dependency as a clinically actionable vulnerability in a defined subset of rhabdoid tumors.

### The core rhabdoid regulatory circuit underlies HGF/MET signaling in ERRTs

Having identified HGF/MET signaling as a selective vulnerability in ERRTs, we next asked how this targetable axis is transcriptionally and epigenetically configured, and whether its activation is linked to the recurrent regulatory programs identified in primary rhabdoid tumors. Consistent with our observations in the patient-derived cell models (**Fig. 5c, d**), MET expression was highest in ERRTs in the single-cell Multiome data, and MET-high tumors also expressed HGF, consistent with an active autocrine HGF/MET signaling axis in ERRT cells (**Fig. 6a; Supplementary Fig. 5a,b**). This transcriptional pattern was mirrored by increased chromatin accessibility at both *HGF* and *MET* loci in ERRTs but not in other rhabdoid samples (**Supplementary Fig. 7a**). To determine whether this pathway was linked to metaprogram-associated regulatory circuitry, we performed patient-level SCENIC+ analyses. Intriguingly, in ERRTs with the highest *HGF* expression, HGF was recurrently linked to the core M12-associated regulators TCF12 and CHD2, together with patient-specific TFs including KLF5, FOXP1 and BACH2 (**Supplementary Fig. 7a,b**). MET showed a similar regulatory architecture, with TCF12 and CHD2 linked to *MET* regulation in six patients, five of whom overlapped with the HGF-high group, alongside additional patient-specific regulators such as KLF5 and HMGA2 (**Fig. 6b; Supplementary Fig. 7a**). This overlapping circuit strongly reinforces HGF/MET axis activation emerging from a shared M12-linked regulatory backbone supplemented by patient-specific inputs, supporting convergent regulatory routes to this targetable signaling dependency. Orthogonal analysis of public DepMap CRISPR/Cas9 screening data (DepMap Public 26Q1; depmap.org) further linked MET expression to MET genetic dependency in ERRTs (**Fig. 6c**). Intriguingly, high MET expression strongly predicted MET dependence in rhabdoid, but not non-rhabdoid childhood cancer models (**Supplementary Fig. 7c**), consistent with MET vulnerability being wired by a core non-redundant rhabdoid regulatory circuitry.

**Fig. 6.**
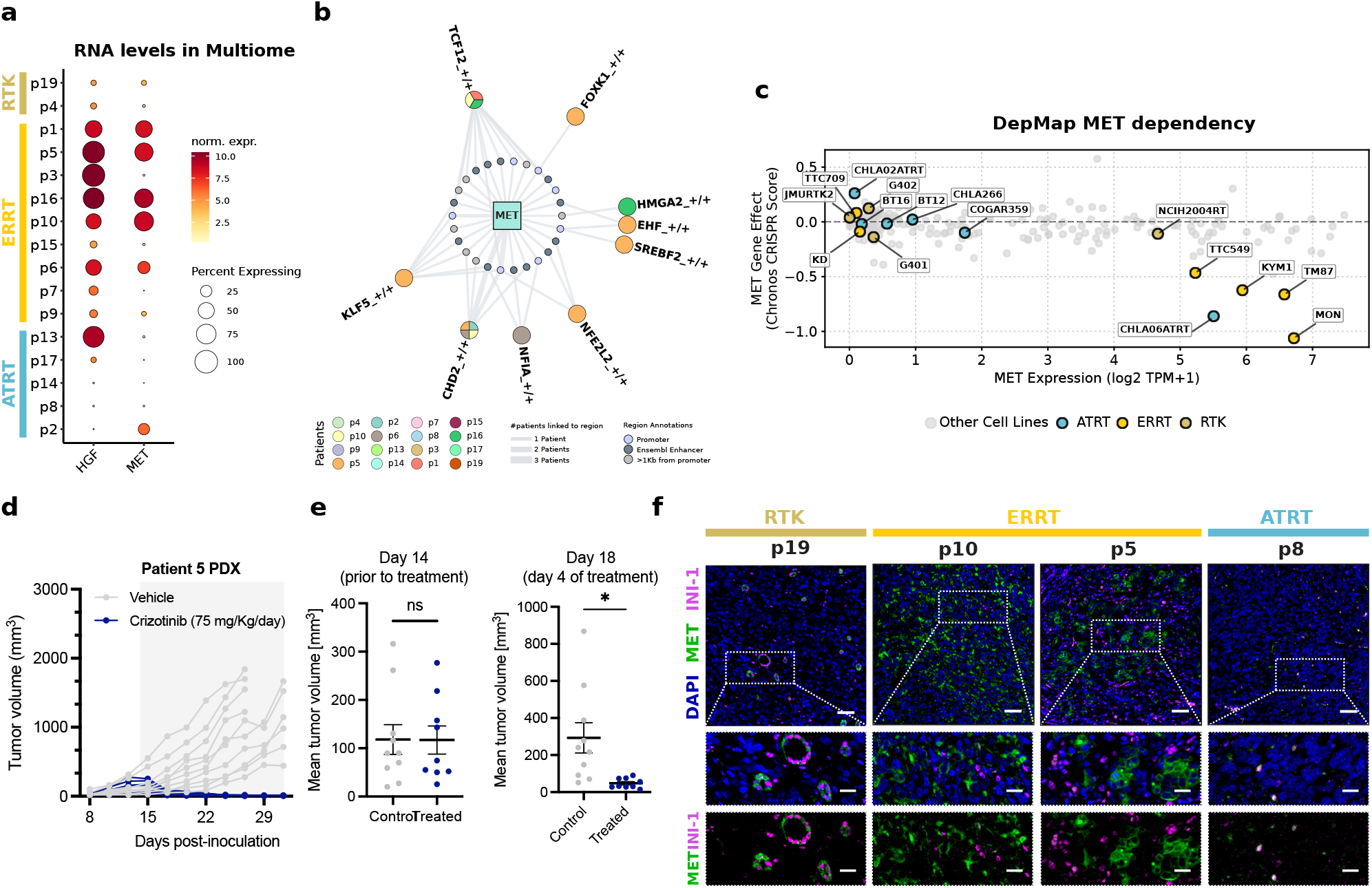
MET expression underlies potent in vivo vulnerability and serves as predictive biomarker for patient stratification. **a**, Dot plot showing *HGF* and *MET* expression within tumor fraction. **b**, Regulatory network showing the patient-specific eRegulons linked to MET. Each eRegulon is depicted as a pie-plot showing in which the patients that eRegulon is linked to MET. Dots representing regulatory regions are classified as *Promoter* if they are within 1Kb of the TSS, *Ensembl enhancers* if they overlap with enhancers in Ensembl regulatory feature annotations, or *>1Kb from Promoter* for all others. Connecting lines width represents the number of patients associated with a specific region. **c**, Scatter plot of *MET Expression (log2 TPM+1)* versus *MET Gene Effect (Chronos CRISPR score)* for 242 cell lines (filtered by *Pediatric type model*) in DepMap (v26Q1). Each point represents a cell line colored as follows: ATRT (blue), RTK (beige), ERRT (yellow), and other cell lines (gray), and all MRT cell lines are correspondingly labeled. Lower Chronos scores represent selective *MET* dependency. **d**, Tumor volume progression over time (days post-inoculation) for vehicle (grey) or 75 mg/kg/day crizotinib (blue) via oral gavage. The grey shaded area indicates the active treatment window starting on day 14 post-inoculation. Data represented are tumor measurements from individual mice. **e**, Tumor volume on days 14 (prior to treatment initiation) and 18 (4 days post treatment). Data represent the mean +/-SEM of individual tumor measurements. Statistical significance was determined using unpaired two-tailed Student‘s *t-*test. **f**, Representative examples of DAPI, MET and INI-1 immunofluorescence on primary tumor specimens (top), and magnifications showing DAPI, MET and INI-1 (middle), or MET and INI-1 only (bottom). Nuclei were counterstained using DAPI. Scale bars, 50 μm (top) and 20 μm (middle and bottom).

### MET inhibition induces marked tumor regression in vivo

We next tested whether MET dependency could be therapeutically exploited in vivo. MET-sensitive patient 5-derived ERRT cells were implanted into NSG mice. Once tumors reached a mean volume of 120 mm^3^, mice were randomized to receive vehicle (n = 10) or the clinically approved MET inhibitor crizotinib at 75 mg/kg (n=9) once daily. Crizotinib elicited a rapid response, with significant antitumor effects as compared to vehicle controls observed as early as 4 days after treatment initiation (**Fig. 6d,e**). Whereas tumors in vehicle-treated animals continued to progress, crizotinib-treated tumors underwent marked regression, with all tumors becoming non-palpable by treatment day 14. Thus, pharmacological inhibition of the MET signalling axis induced a rapid and deep response in ERRT PDX models, supporting the clinical translation of MET-directed therapy in ERRT.

Because clinical implementation of MET-targeted therapy requires a readily applicable patient selection strategy, we next assessed whether MET protein abundance in patient tumors could serve as a stratification biomarker. Immunohistochemical analysis revealed strong membrane-localized MET staining in ERRTs, whereas RTKs and ATRTs were largely negative (**Fig. 6f; Supplementary Fig. 7d**). In addition, co-staining for SMARCB1 (INI-1) further showed that MET expression colocalized to SMARCB1-deficient tumor cells in ERRTs, confirming MET expression specifically within the malignant ERRT compartment. Thus, our findings support tumor-cell MET expression as a candidate stratification biomarker identifying MET-dependent rhabdoid tumors.

Collectively, by integrating single-cell regulatory architecture, spatially-resolved signaling and functional perturbation, we uncover a recurrent yet subtype-enriched HGF/MET dependency in ERRTs and nominate MET expression as a biomarker for therapeutic patient stratification.

## Discussion

Malignant rhabdoid tumors are deadly childhood cancers that are characterized by the unifying event of SWI/SNF complex dysregulation – most commonly through functional inactivation of SMARCB1 - on an otherwise mutationally quiet background ^(1,2,6,9)^. Despite this apparent genomic simplicity, diverse anatomic presentation and distinct molecular subtypes ^(2,12–14)^ hint at a heterogeneous disease. Mapping rhabdoid tumor heterogeneity both within and across samples offers the opportunity to unearth both unifying and distinguishing features, enabling the understanding of shared biology, as well as the identification of patient subgroups who may benefit from highly specific targeted therapies.

Here we combined single-cell Multiome profiling with spatial transcriptomics across a rhabdoid tumor cohort that spans both molecular and anatomical diversity. Malignant cell populations displayed striking intra- and inter-tumor heterogeneity. Within-tumor variation in transcriptional states was, in part, spatially patterned and associated with neighborhood context. Despite inter-sample variability, we identified recurrent malignant transcriptional programs shared across patients and found that these metaprograms were associated with a *TCF12*/*ZNF148*/*CHD2*-centered core regulatory circuit (CRC). Thus, our data extends previous bulk-level analyses ^(12–15)^ by providing single-cell and spatial resolution, and complements a recent ATRT-focused single-cell atlas ^(21)^ by expanding multimodal and spatial resolution to diverse extracranial presentations of rhabdoid tumors, thus enabling broader insights into rhabdoid cell biology. Indeed, leveraging our multimodal data, we identify and functionally validate HGF/MET signaling as a clinically actionable vulnerability in a defined subset of rhabdoid tumors, with MET expression emerging as a biomarker for patient stratification.

In the context of genomic simplicity that characterizes SMARCB1-deficient rhabdoid tumors ^(6,9)^, this heterogeneity argues for a prominent contribution of non-genic mechanisms of transcriptional diversification, rather than capturing transcriptional profiles of genetically distinct malignant cell clones. Such mechanisms can arise cell-intrinsically, for instance through stochastic state fluctuations ^(43,44)^, developmental-like differentiation hierarchies previously described in childhood cancers ^(21,45)^, or can be imposed or reinforced by microenvironmental cues or tumor-stroma interactions ^(46)^. In our cohort, the observation that distinct malignant transcriptional states are spatially patterned, with certain transcriptional states concentrated in tumor dominated areas, or colocalizing with stromally enriched (often non-contiguous) regions, supports a prominent role for microenvironmental cues in shaping intratumor heterogeneity and cell state plasticity.

Considering this intratumor transcriptional diversity, it is perhaps not surprising that tumors that arise in anatomically diverse locations show substantial inter-patient molecular divergence, potentially reflecting long-term co-adaptation to distinct tissue niches. By focusing on co-expressed gene networks, we identified recurrent transcriptional metaprograms shared across patients. This program-centric strategy mirrors prior work showing that highly heterogeneous malignant compartments can be decomposed into a defined set of recurrent expression programs across tumors ^(47,48)^. The 19 metaprograms we identified captured conserved features of rhabdoid biology present across anatomical sites and molecular subtypes, alongside programs that were subtype-skewed or coupled to microenvironmental signaling. A central finding of our work was a ‘core’ rhabdoid metaprogram (M12), shared across most tumors and enriched for pathways consistent with known consequences of SMARCB1 loss, including chromatin regulation and stress adaptation programs linked to proteostasis and unfolded protein response. Notably, M12 was variably active between subpopulations of the same tumor, indicating that shared rhabdoid biology co-exists with marked cell state diversity rather than being uniformly expressed throughout each tumor. Consistent with this, we observed spatial segregation of M12 core and M18 edge-associated programs, supporting niche-imposed state changes and suggestive of transcriptional, and potentially phenotypic, plasticity in rhabdoid tumor cells. Notably, phenotypic plasticity can enhance cellular fitness under stress, for instance by enabling acquired resistance to therapy - an important clinical challenge in the treatment of rhabdoid tumors ^(49)^.

To connect the identified transcriptional metaprograms to upstream regulatory circuits, we inferred eRegulons associated with each programme, providing a mechanistic link between co-expressed gene networks and candidate transcription factor drivers. Interestingly, the M12 core rhabdoid metaprogram was associated with a compact set of eRegulons centred on *TCF12, ZNF148* and *CHD2*, consistent with shared underlying regulatory wiring across tumors and supporting this set as a CRC downstream of SMARCB1 loss. Moreover, we identified that the M2 metaprogram was in part driven by *OTX2*, a key transcriptional regulator involved in anterior neuroectoderm patterning and neurogenesis, and consistent with promotion of the ATRT-like developmental programs enriched in M2 ^(50)^.

Taken together, we propose a modular framework in which the malignant phenotype is defined by factors related to the initiating genetic event (SMARCB1 loss), the cell’s epigenetic state (potentially linked to the state in which transformation occurred), and environmental inputs. In this view, a shared core rhabdoid program - defined by the activation of a rhabdoid-specific CRC - is layered with subtype-skewed programs and further diversified over time by niche-imposed co-adaptation, producing the transcriptional heterogeneity we observe across samples.

Our principal translational finding is the identification of HGF/MET signaling as a selective and clinically actionable dependency in a defined subset of rhabdoid tumors. In contrast to other therapeutic intervention axes nominated by our multi-omic analyses, which displayed either limited efficacy (WNT-pathway/MDK) or limited tumor selectivity (chemotherapeutic, epigenetic or metabolic inhibitors), MET inhibition produced robust and tumor-selective cytotoxicity in ERRTs. Multiple orthogonal lines of evidence support this dependency: (1) HGF/MET signaling axis was identified in a subset of patient tumors; (2) patient-derived ERRT models displayed exquisite sensitivity to pharmacologically distinct MET inhibitors, but not AXL inhibition; and (3) sensitive but not resistant models showed HGF-dependent MET pathway activation. Importantly, external CRISPR/Cas9 dependency data from DepMap independently linked MET expression to MET genetic dependency in rhabdoid tumor models, but not broadly across other MET-expressing pediatric cancer models. This points to a distinctive feature of MET-positive rhabdoid tumors: HGF/MET signaling appears to be embedded within the regulatory logic of the malignant state, rather than functioning as an isolated pathway activation event. Consistent with this model, *HGF* and *MET* expression were linked to the M12-associated TCF12 and CHD2 eRegulons, together with additional tumor-specific co-regulators, suggesting that a shared rhabdoid regulatory backbone can be coupled to patient-specific inputs to activate an autocrine HGF/MET program. This convergence may explain why MET expression marks dependency in rhabdoid tumors but not in other pediatric cancers, nominating HGF/MET signaling as an identity-associated and therapeutically tractable vulnerability.

From a clinical perspective, MET is an attractive target, since selective MET inhibitors are already FDA/EMA approved and widely used in biomarker-defined settings, most prominently for advanced non-small cell lung cancer (NSCLC) with MET exon 14 skipping, where MET-targeted tyrosine kinase inhibitors (TKIs) have shown clinically meaningful responses ^(51,52)^(NCT02414139; NCT02864992). Importantly, the clinical application of MET-directed therapies is expanding beyond genotype-defined MET alterations to include protein-based biomarkers, with MET overexpression by IHC used to select patients for MET-directed treatment strategies ^(53)^(NCT03539536), establishing a clear precedent for IHC-guided patient stratification for MET-targeted treatments. Of note, the MET targeting landscape is evolving beyond small-molecule MET inhibition towards antibody-based modalities, including MET-directed antibody-drug-conjugates (ADCs)^(53)^, ligand/receptor-directed antibodies (e.g. anti-HGF antibody ficlatuzumab; NCT03422536)^(54)^, and bispecific formats that incorporate MET as a therapeutic handle (NCT04868877). Specific design features of these next-generation treatments may be advantageous in heterogeneous and spatially structured cancers such as rhabdoid tumors: for instance, ADCs can mediate bystander killing through payload diffusion, potentially extending drug efficacy beyond target-high cells ^(55)^. Moreover, paracrine HGF/MET signaling is an established driver of invasive growth programs and is broadly implicated in tumor progression ^(56,57)^, providing a clear rationale for ligand-neutralizing strategies in settings where stromally-derived HGF may contribute to pathway activation.

In conclusion, our data indicate that MET IHC is a clinically feasible strategy to identify the subset of rhabdoid tumors most likely to benefit from MET inhibition, providing a practical route to biomarker-informed clinical deployment. Conceptually, MET IHC could be smoothly incorporated into pediatric umbrella screening frameworks such as Pediatric MATCH (NCT03155620) to enable prospective selection of patients for MET-directed treatments, and thus opening a clear path to biomarker-selected MET targeting in rhabdoid tumors or even in MET-expressing childhood cancers more broadly. Together, our work provides a molecularly grounded and compelling opportunity to bring MET-directed precision therapy to MET-expressing childhood cancer patients in the near future.

### Limitations of the study

This study has several limitations. First, the extreme rarity of rhabdoid tumors constrained cohort size, limiting fine-grained subtype-specific comparisons. Nonetheless, our study offsets this limitation through deep molecular profiling, thoughtful data integration and ultimately functional validation in matched patient-derived models. Importantly, although limited in size, our cohort spans substantial anatomical and molecular diversity and, to our knowledge, represents the most comprehensive multimodal MRT dataset assembled to date. Second, our spatial analyses do not provide direct single-cell-resolution measurements, but instead rely on spot-based transcriptomics coupled with deconvolution and computational inference. Nonetheless, this framework enabled us to identify recurrent spatial neighborhoods and robust, recurrent cell-cell communication programs at the cohort level. Finally, while MET dependence is supported by orthogonal analyses in primary tumors and patient-derived models, additional validation in independent cohorts and prospective translational settings will be important to define the full scope of biomarker-guided MET targeting in rhabdoid tumors.

## Resource availability

### Lead contact

Further information and requests for resources and reagents should be directed to and will be fulfilled by the Lead Contact, Alexandra Avgustinova.

### Materials availability

Patient-derived models and other materials generated in this study are available from the Lead Contact upon reasonable request. Distribution may require completion of a material transfer agreement and will be subject to institutional policies and any applicable ethical or legal restrictions associated with human-derived materials.

### Data availability

The raw data (FASTQ files) for single-cell Multiome and Visium 10x Genomics data are deposited in the European Genome-phenome Archive (EGA code EGAS50000002211) (https://ega-archive.org/studies/EGAS50000002211). The standardized processed data objects are on Zenodo under request (https://doi.org/10.5281/zenodo.19666296).

These include single-cell Seurat objects (.rds) and spatial transcriptomics AnnData objects (.h5ad) These objects contain the integrated and annotated expression matrices used in the analyses and visualizations presented in this study. MET expression and gene effect scores (Chronos) can be consulted at DepMap (v26Q1) (https://depmap.org/portal/).

### Code availability

Computational code and analysis scripts used to generate the results are available in the following GitHub repository at: https://github.com/Single-Cell-Genomics-Group-CNAG-CRG/Multimodal-Analysis-Rhabdoid-Tumors

## Acknowledgments

AA acknowledges funding from the European Research Council (ERC-StG 101076506) and the Ramón y Cajal fellowship (RYC2019-027738-I). This project was supported by the PID2020-118241RA-I00 project, and an FPI PhD fellowship contract for ACX (PRE2021-098532), both funded by the Spanish State Research Agency (AEI). This work has also been supported by the XXV FERO award (Fundación FERO) granted to AA. HH received funding from the Ministerio de Ciencia e Innovación, Proyectos I+D Generación de Conocimiento 2020 (PID2020-115439GB-I00), the Obra Social Fundación La Caixa (CAIXARESEARCH health call 2022, HR22-00172) and the Agencia Estatal de Investigación, Lineas estratégicas (PLEC2021-007654). IS work is supported by Juan de la Cierva fellowship (FJC2021-047130-I) funded by the MCIN and Spanish State Agency of Investigation (AEI) (10.13039/501100011033) and also the European Union’s NextGenerationEU/PRTR fund project. QR work was supported by the Scientific Foundation of the Spanish Association against Cancer (TALEN258116ROVI) and the European Union through the AECC Talent Programme (GA101081298). We thank the *Market Solidario* and *Pulseras Candela* for their continued support. We acknowledge the support of the Histopathology and Biostatistics/Bioinformatics Facilities, and the Drug Screening Platform of the Institute for Research in Biomedicine Barcelona, and the Anatomical Pathology Department of Sant Joan de Déu Children’s Hospital Barcelona, for supporting our work. We are indebted to the Biobanc de l’Hospital Infantil Sant Joan de Déu per a la Investigació (BISJDI), integrated in the Spanish Biobank Network of ISCIII, for the sample and data procurement. We thank Ireen J. Kal for her assistance and generation of preliminary regulon data. Finally, we extend our deepest gratitude to the patients and their guardians, whose participation made this study possible.

## Author contributions

AA and HH conceived and supervised the project. IS performed the processing of sample data for Multiome and Visium assays and performed all the spatial transcriptomic analysis. ACX established all patient-derived cell lines and performed functional validation assays. QR did all the gene regulatory analysis. EMA helped with the functional validation assays. IS, ACX, QR, EMA did all the figures. AA, IS, ACX, QR and EMA interpreted the results and wrote the manuscript. SM, JCN, HH and DaM participated in providing key additional insights and discussion. CJRH coordinated all the sample acquisition and storage. SM ran hdWGCNA and provided output results and co-expression network plots. AdC computed RT signature by analyzing public RNA seq data. IR annotated H&E tissue slides. AUB and AO both help in establishing the patient-derived tumor cell lines and provide further support to experiments. MC and IR performed the drug screening. SGP and ERN performed the immunohistochemistries. DoM and GC generated the Multiome 10X Genomics dataset. PL and MR generated the Visium 10X Genomics dataset under the supervision of APR. CEL, AlC and JM are responsible for the storage of the samples, diagnosis of the tumors and providing clinical information.

## Methods

### Ethical compliance

All research activities comply with ethical regulations and were performed in accordance with the Declaration of Helsinki. Patient material was procured and processed through the Sant Joan de Déu Barcelona Children’s Hospital BioBank for Research (BHISJDI), integrated in the Spanish Biobank Network of Carlos III Health Institute (ISCIII), after written, signed, and dated informed consent from the patients’ legal guardians. Access to the samples and the study protocol were reviewed and approved by the Hospital Sant Joan de Déu Drug Research Ethics Committee (CEIm) under approval number PIC-30-21. All human rhabdoid tumor biopsies safeguarded in the BHISJDI were managed in strict accordance with the Spanish Biomedical Research Law (Law 14/2007) and Royal Decree 1766/2011.

### Patient specimens and biobanking

Fresh tumor samples collected between 2007 and 2024 were gently minced into small fragments (0.5 – 2 mm^3^) using sterile scalpels and washed in pre-warmed RPMI 1640 medium (Gibco, 31870025). Following centrifugation at 200 × g for 5 min, fragments were resuspended in either fetal bovine serum (FBS) containing 10 % dimethyl sulfoxide (DMSO; PanReac AppliChem, A3672), or Synth-a-Freeze cryopreservation medium (Gibco, A1254201) for subsequent cryopreservation. To ensure optimal cellular viability, samples underwent gradual cryopreservation at −80°C for 48 hours before being transferred to liquid nitrogen tanks for long-term storage. Additionally, tumor fragments (0.5 - 2 mm^3^) were snap-frozen in liquid nitrogen and stored at −80°C by the BHISJDI for future molecular analysis.

### Clinical diagnosis and molecular MRT classification

Histopathological confirmation of INI-1 (SMARCB1) negative was conducted by the Pathology Department at Hospital Sant Joan de Déu Children’s Hospital (HSJD) through immunohistochemical (IHC) staining. For rhabdoid tumor molecular stratification, primary tumor classification was performed by the Molecular Diagnostics Laboratory based on DNA methylation profiling (Infinium HumanMethylation450 or MethylationEPIC BeadChip), and raw methylation data was processed using the Epignostix Heidelberg Classifier (https://app.epignostix.com) for central nervous system or sarcomas to determine the specific MRT subgroup.

To assess *SMARCB1* genomic alterations in both primary tumors and patient-derived cell lines, we performed Multiplex Ligation-dependent Probe Amplification (MLPA) using the SALSA MLPA Probemix for SMARCB1 (MRC Holland, P258-025R) according to the manufacturer’s instructions. Data analysis was performed using Coffalyser software v40129.1959 (MRC Holland, Netherlands). Small genomic alterations, specifically short indels and point mutations, were confirmed via Sanger sequencing within the *SMARCB1* coding regions.

### Immunohistochemistry/Immunofluorescence staining

Fresh tumor biopsies were obtained from resected MRT tumors, routinely fixed in 10 % neutral buffered formalin, paraffin-embedded and blocks were cut into 3 μm sections and deparaffined at 60°C for 45 min. For antigen retrieval, sections were either boiled in Tris-EDTA pH 9 solution or autoclaved at 1 atm of pressure in citrate buffer pH 6 for 20 minutes. Immunohistochemistry stainings were either conducted in a Leica Bond-Prime (Leica, USA) or Dako Autostainer Link 48 (Dako, USA) systems, except for MET and HGF that were manually stained. Primary patient samples were characterized by immunohistochemical detection of INI-1 (1:200; BD Biosciences, 612110), HGF (1:100; Thermo, PA5-115354) and MET (1:50; CST, 8198), and counterstained with hematoxylin and eosin (H&E). Stained immunohistochemistry sections of MRT tumor lesions were digitalized in a high-resolution NanoZoomer 2.0 HT (Hamamatsu Photonics, Japan) and visualized using the NDP.view2 Image software U12388-01 (Hamamatsu Photonics, Japan). For immunofluorescence stainings, 3 μm sections were incubated with the primary antibodies INI-1 (1:200; BD Biosciences, 612110) and MET (1:50; CST, 8198). Nuclei were counterstained with 1 μg/mL DAPI. Stained sections were scanned using the x20 objective in a PhenoImager HT 2.0 (Akoya Biosciences/Quanterix, USA). Fluorescent images were visualized in QuPath 0.7.0 (University of Edinburgh, Scotland) and further processed in FIJI ImageJ software 1.54p (National Institutes of Health, USA).

### Sample preparation for 10X Genomics v1 3’ Visium Spatial Gene Expression

Tumors were resected, embedded in OCT embedding matrix and snap frozen. 8-10 sections of 50 µm were cut in the region proximate to the tissue section used for Spatial Transcriptomics, by using a cryostat (Leica CM1950).

Frozen tissue samples were tested for RNA quality with RIN > 7.0 (RNA pico, Agilent) and a tissue optimization experiment (10x Genomics, Visium Spatial Tissue Optimization, Rev A) was performed with imaging of fluorescence footprint on upright epifluorescence (Nikon Eclipse E800) identifying 3 min as optimum permeabilization time.

The samples were then processed for full ST experiment as per manufacturer’s instructions (10x Genomics, Visium Spatial, Rev D) being cut in a pre-cooled cryostat at 10 μM thickness onto four 6.5mm × 6.5mm capture areas with 5000 oligo-barcoded spots. Slides then underwent fixation and H&E staining with immediate imaging on NanoZoomer S60 (Hamamatsu). Tissue underwent permeabilization with proprietary enzyme (3 min), reverse transcription and second strand synthesis performed on the slide with cDNA quantification undertaken with qRT-PCR using KAPA SYBR FAST-qPCR kit (KAPA Biosystems) and analyzed on the LightCycler®480 Real-Time PCR System (Roche). qRT-PCR results (Cq value at 25 % of peak fluorescence) informed cDNA amplification. Following library construction as per manufacturer’s instructions ST libraries were quantified by High Sensitivity DNA Assay 2100 Agilent. Libraries were sequenced on NovaSeq6000 platform (Illumina, US) using 100 base-pair paired-end dual-indexed set up. Around 200 million pair-ended reads were obtained for each tissue section. Read 1, 28 cycles, i7 index 10 cycles, i5 index 10 cycles and Read 2 were sequenced with 82 cycles respectively.

### Sample preparation for 10X Genomics Single Cell Multiome ATAC + Gene Expression

Nuclei isolation was performed following the “Nuclei Isolation from Complex Tissues for Single Cell Multiome ATAC + Gene Expression Sequencing’’ demonstrated protocol (10x Genomics; CG000375). After a first tissue lysis step with NP40 Lysis Buffer, a minimum of 200,000 nuclei were FACS sorted upon staining with 1 µg/ml DAPI (Merck) using a FACSAria™ Fusion Flow Cytometer (BD Biosciences). This was done in order to eliminate tissue debris and ambient DNA/RNA. Sorted nuclei were then permeabilized with 0.1X Lysis Buffer, washed and resuspended in the appropriate volume of 1X NB buffer (1 M MgCl2, 1 M Tris-Cl (pH 9.5), 5 M NaCl, 20 % Tween 20) supplemented with 1U/µl of RNAse inhibitors (PN. 3335402001, Sigma). After filtration with 40 μm Flowmi Cell Strainer (PN. BAH136800040-50EA, Merk), nuclei were stained with Trypan blue and counted manually using a Neubauer chamber. Gene expression (GEX) and ATAC-seq libraries were prepared following the Chromium Next GEM Single Cell Multiome ATAC + Gene Expression User Guide (10x Genomics; CG000338). Transposed nuclei were partitioned into GEMs (Gel Beads in Emulsion) using the Chromium Controller with Chip J aiming at a target recovery of 6000-10.000 nuclei per sample, depending on the final number of nuclei obtained. After GEMs incubation for mRNA reverse transcription and transposed DNA barcoding, the resulting cDNA and barcoded gDNA were purified and pre-amplified during 7 cycles, following the 10x Genomics protocol. The sample was split into 35 µl for cDNA amplification and 40 µl for ATAC library construction. After a clean-up, 35 µl of the pre-amplified cDNA were amplified with 7 additional PCR cycles. The resulting cDNA was quantified on an Agilent Bioanalyzer High Sensitivity chip (Agilent Technologies) and 100 ng were used for GEX library preparation. GEX libraries were indexed with 13 cycles of amplification using the Dual Index Plate TT Set A (10x Genomics; PN-3000431). In parallel, 40 µl of the pre-amplified DNA were indexed with 7 cycles of amplification using the Sample Index N Set A (10X Genomics; PN 3000427). Size distribution and concentration of full-length GEX and ATAC-seq libraries were verified on an Agilent Bioanalyzer High Sensitivity chip (Agilent Technologies). Finally, sequencing of GEX libraries was carried out on a NovaSeq 6000 sequencer (Illumina) using the following sequencing conditions: 28 bp (Read 1) + 8 bp (i7 index) + 0 bp (i5 index) + 89 bp (Read 2), to obtain approximately >20,000 paired-end reads per cell. ATAC-seq libraries were also sequenced with a NovaSeq 6000 sequencer (Illumina) using the following conditions: 50 bp (Read 1N) + 8 bp (i7 Index) + 16 bp (i5 Index) + 49 bp (Read 2N), aiming at a sequencing depth of >25,000 reads/nucleus.

### 10X Genomics Single Cell Multiome Analysis

FASTQ files from Single Cell Multiome libraries (RNA and ATAC) were processed using Cell Ranger ARC (v2.0.0) pipeline that jointly maps gene expression and chromatin accessibility to the human GRCh38 reference (GENCODE v32). We performed data analysis using R (v4.1.1), Seurat (v4.0.5)^(58)^ and Signac (v1.4.0)^(59)^. A detailed list of software dependencies and specific package versions for each computational step is documented in the R notebooks and HTML reports available on the project’s GitHub repository.

#### Quality Control (QC)

Multi-omic Seurat objects were generated from raw Cell Ranger ARC count matrix outputs for each patient. For the ATAC-seq assay, we filtered peaks to exclude non-standard chromosomes and regions in the hg38 unified blacklist curated by Signac developers. RNA and ATAC data were then integrated into a single object, retaining only shared barcodes to ensure consistent multi-omic profiling. To facilitate quality control, we used Signac to calculate fragment-level statistics, including the Fraction of Reads in Peaks (FRiP), nucleosome signal, and TSS enrichment.

We applied a permissive filtering strategy to preserve biological diversity for downstream, cluster-specific cleaning. RNA filters included a library size of 400-45,000 UMIs, 200-10,000 detected genes, and maximum thresholds of 20 % mitochondrial and 60 % ribosomal content. ATAC filters required 1,000-25,000 fragments in peak regions, a FRiP score > 0.15, a TSS enrichment score > 1, and a nucleosome signal < 2.

Potential doublets were identified using modality-specific computational tools: Scrublet (v0.1.0)^(60)^ for the RNA assay and the addDoubletScores function from ArchR (v1.0.2)^(61)^ for the ATAC data. The resulting doublet scores were stored within the Seurat object metadata for downstream evaluation. In short, to avoid the arbitrary loss of valid biological states, we postponed final doublet exclusion to after an initial round of unsupervised clustering (explained in the following sections). This strategy allowed us to validate doublet predictions by identifying clusters that exhibited both high computational scores and mixed marker expression profiles.

#### Processing of samples and first clustering

RNA counts were normalized by dividing the raw counts of each cell by its total counts, followed by a scale factor of 10,000 and a log(1+x) transformation. Simultaneously, ATAC peak counts were normalized using the Term Frequency-Inverse Document Frequency (TF-IDF) method. In addition, we computed the cell cycle score using the gene list provided by the function *cc*.*genes* and added cycle ‘phases’ (G1, G2M, and S) to the metadata to account for proliferating cells in downstream analysis. To reduce noise and highlight biological signals, we selected highly variable genes for the RNA modality. This gene selection was performed modulating the mean and the variance of the log-normalized values using *modelGeneVar* function from scran (v1.22.1) and setting an FDR threshold of 0.05 at *getTopHVGs* function to obtain a list of highly variable genes (HVGs) These values were z-score transformed and subjected to Principal Component Analysis (PCA). For the ATAC modality, we identified the top peaks based on total counts and performed dimensionality reduction using Singular Value Decomposition (SVD), as it is better suited for the sparsity of ATAC data than standard PCA, providing what is known as latent semantic indexing (LSI). For RNA data, principal components (PC) were selected for each sample based on the Elbow Plot of the standard deviation. For ATAC data, we discarded the first LSI component in all patients as we checked it was correlating with sequencing depth.

To integrate the two modalities, we constructed a Weighted Nearest Neighbor (WNN) graph for each patient. This unsupervised framework allowed for the joint analysis of RNA and ATAC data to define neighbors. Uniform Manifold Approximation and Projection (UMAP) was used to visualize data. Unsupervised clustering was then performed using the Louvain algorithm, with the resolution chosen for each patient to ensure the accurate capture of tumor heterogeneity and major cell populations. Finally, we identified cluster-specific marker genes using the Wilcoxon Rank Sum test. These top differentially expressed genes guided the initial annotation of cells into broad categories, including tumor cells, immune cells, and other stromal components. At this point, we already observed low quality cells clustering in p5 and p9 and removed them.

#### RNA ambient removal

During the initial unsupervised clustering, we observed evidence of ambient RNA contamination, which manifested as the cross-expression of lineage-specific markers between distinct cell populations. This was particularly evident in several patients where the pan-immune marker *PTPRC* (CD45) was detected within malignant cell clusters, and tumor-specific transcripts were found at low levels in non-tumor cells. To mitigate these artifacts, we applied DecontX (v1.2.0) to estimate and remove the ambient RNA signal from the RNA assay. We run DecontX using the filtered count matrix and initial cluster labels as inputs. To accurately model the background “soup,” the original raw feature-barcode matrix from Cell Ranger was provided as a reference for ambient RNA distribution. The algorithm further used highly variable genes to refine the estimation of native versus contaminating transcripts. This approach allowed for the calculation of a contamination score per cell and the generation of a decontaminated “native” expression matrix, which was used for all subsequent transcriptomic analyses.

To ensure that downstream dimensionality reduction and clustering were driven by robust biological signals rather than outlier transcripts, we performed a per-patient evaluation of total gene counts. For each sample, we visualized the genomic distribution of total pseudo-bulk transcripts using a Manhattan plot, where genes were ordered by genomic coordinates and plotted against their cumulative counts. This approach allowed for the identification of genes with disproportionately high expression levels (such as MALAT1) which exceeded a count threshold (e.g., 150,000 counts). To prevent these dominant features from masking more subtle biological variation, genes exceeding this threshold were flagged for each patient and systematically excluded from the set of highly variable genes (HVGs) used for subsequent integration and PCA.

#### Annotation of patient-derived Multiome clusters

Following ambient RNA decontamination and the exclusion of dominant transcripts, the refined expression matrices were processed individually. For the transcriptomic assay, counts were normalized using a global-scaling method that divided feature counts by the total counts per cell, multiplied by a scale factor of 10,000, and log-transformed. Feature selection was performed by identifying HVGs based on their mean-variance relationship. The data were then scaled, and Principal Component Analysis (PCA) was performed, with the optimal number of dimensions determined for each patient via elbow plots.

For the chromatin accessibility assay, dimensionality reduction was conducted using Latent Semantic Indexing (LSI). To integrate the two modalities, we computed a Weighted Nearest Neighbor (WNN) graph, which served as the basis for UMAP visualization and unsupervised Louvain clustering.

Clustering resolutions were adjusted on a per-patient basis to ensure the captured variability reflected true biological heterogeneity rather than technical noise. By analyzing dimensionality reduction loadings and lineage-specific markers (**Supplementary Fig. 1, Extended Data Fig. 1**), we effectively partitioned the tumor microenvironment (TME). Malignant cells were distinguished from stromal and immune populations – including macrophages, T cells, endothelial cells, and fibroblasts – through the inspection of canonical markers (e.g., PTPRC for immune cells, PECAM1 for endothelial cells, and several collagenases for fibroblasts). Malignant clusters were further characterized and annotated based on the expression of their top differentially expressed genes (see **Supplementary Fig. 1, Extended Data Fig. 1**).

#### Integration of samples

To perform a comparative analysis across the entire cohort, individual Seurat objects were merged into a single integrated object. To resolve differences in peak calling between samples, we used the UnifyPeaks() function from the Signac package to create a common set of peaks across all datasets. The chromatin accessibility signal for each sample was then re-quantified using this unified peak set and the original fragment files to generate a standardized counts matrix.

A new Chromatin Assay was initialized for the merged object using these unified peak counts, and the corresponding RNA assay matrices were concatenated. To reduce computational overhead we removed lowly expressed genes (less than 5 counts). Finally, the fragment files from all patients were merged into a single master file, with cell barcodes uniquely prepended by their respective patient IDs to prevent barcode collisions across the cohort. This step was critical to enable efficient cross-sample interrogation of the raw data, allowing for the generation of aggregated genomic coverage tracks and peak visualizations across the integrated cohort.

The unified cohort object underwent processing. The gene expression assay was log-normalized and we normalized the common peaks with TF-IDF. To further mitigate batch-specific biases and gender-related confounding effects, sex-linked (dimorphic) genes (62) were removed from the matrices prior to feature selection as well as the previously identified dominant outlier genes were also excluded. To ensure the selection of robust biological markers across sequencing subprojects, we performed batch-aware feature selection using scran. In short, we modeled the mean-variance relationship within each subproject independently, identifying significantly variable genes using a FDR threshold of 0.05. To prioritize genes with high cross-batch reproducibility, only those identified as highly variable in at least two independent subprojects were retained for downstream dimensionality reduction and integration. For ATAC peaks, we applied a frequency-based filter with a minimum cutoff of 10. This step restricted the analysis to peaks with at least ten total counts across the dataset, thereby excluding sparse low-frequency regions.

To evaluate the drivers of variation in the integrated cohort, we performed a Principal Component Regression (PCR) using metadata covariates. This analysis revealed a high correlation between the first several Principal Components (PCs) and the patient identity, indicating a strong batch effect. To mitigate this, we applied the Harmony integration algorithm to both the RNA (PCA) and ATAC (LSI) embeddings. These harmonized components were then utilized to construct a WNN graph for joint-modality visualization and clustering. Initial unsupervised clustering using the Louvain algorithm identified a cluster (Cluster 0) significantly enriched for ribosomal genes. Despite previous quality control, patient-specific variation in ribosomal content continued to drive the integration. To focus the analysis on relevant biological variation, we performed a second iteration of the processing pipeline with expanded gene exclusion criteria. That is, all ribosomal genes were removed from the count matrix and mitochondrial genes were excluded from HVG together with outlier high count genes and sex-linked dimorphic genes (as before) to prevent them from contributing to PCA loadings.

The refined object was re-integrated and clustered across a range of resolutions. At a lower resolution (0.05), clusters still remained largely patient-specific (see **Fig. 2b**); PCR analysis of the harmonized PCs confirmed that while PC1 captured primary biological lineages, subsequent components remained highly correlated with patient ID, reflecting the high inter-patient heterogeneity characteristic of malignant cells (see **Supplementary Fig. 3e,f**).

Notably, two populations (Cluster 0 and Cluster 1) showed more inter-mixing across patients. Over-Representation Analysis (ORA) using EnrichR (v3.2) and several databases (MSigDB_Hallmark_2020, KEGG_2021_Human, Reactome_2022, WikiPathways_2024_Human) on the differentially expressed (DE) genes of these clusters revealed enrichment for Oxidative Phosphorylation (Cluster 0) and pathways related to cilia (Cluster 1), the latter of which has already been described in ATRTs ^(21)^(**Supplementary Fig. 3c,d, Extended Data Fig. 3a**).

#### Clustering and annotation of immune cells

To achieve a higher resolution of the immune landscape, we performed a targeted sub-analysis of the immune compartment. Immune cells were subsetted from the integrated multiomic object and re-processed independently. Following subsetting, genes with zero expression across the subset were removed, and the raw counts were re-normalized using log-normalization as previously described.

HVGs were identified using a patient-matched modeling approach. We utilized scran to model the mean-variance relationship, specifically excluding mitochondrial genes, sex-dimorphic genes, and genes with aberrantly high counts to prevent technical or biological biases from driving the embedding. Genes with a FDR < 0.05 in at least two patients were included in the final consensus HVG list. This feature set was scaled, and the first 30 PCs were selected for downstream analysis based on elbow plot heuristics.

To leverage established transcriptomic markers for cell-type identification, this sub-analysis focused exclusively on the RNA modality. We employed Harmony to mitigate inter-patient batch effects, followed by the construction of a K-nearest neighbor (KNN) graph. Initial unsupervised clustering was performed using the Louvain algorithm (resolution = 0.3).

Through iterative marker gene inspection, we identified and removed latent doublet clusters that were not apparent in the global analysis. Targeted sub-clustering was also performed on cluster 4 to resolve distinct lineages. The final, “clean” immune dataset was re-processed using the same pipeline with a final clustering resolution of 0.2 and specific sub-clustering of cluster 6, resulting in the final immune populations represented in **Supplementary Fig. 5c-e**.

#### Clustering and annotation of other stromal cells

To further resolve the complexity of the non-malignant stroma, we performed a sub-analysis of the non-immune stromal compartment. Stromal cells were subsetted from the global integrated object and subjected to an iterative refinement pipeline. Initial processing involved the removal of unexpressed genes and log-normalization.

HVGs were identified using a patient-specific modeling approach with scran. To ensure the embedding was driven by robust biological signals, we excluded mitochondrial genes, sex-dimorphic genes, and genes with aberrantly high counts. We defined a consensus HVG list comprising genes that reached significance FDR < 0.05 in at least two patients. This feature set was scaled, and the first 25 PCs were retained for downstream analysis based on variance stabilization.

Following the same rationale as the immune sub-analysis, we utilized the RNA modality for high-resolution lineage identification. Patient-level batch effects were mitigated using Harmony, followed by the construction of a KNN graph and Louvain clustering (resolution = 0.3).

During the initial clustering pass, cluster 4 was identified as a low-quality/mixed population and was subsequently removed. The remaining “clean” stromal dataset was re-integrated and re-clustered at a resolution of 0.3. To achieve the final granularity required for annotation, we performed targeted sub-clustering of cluster 4. This iterative approach enabled the final annotation of stromal sub-populations (e.g., mostly fibroblast and endothelial cells) as detailed in **Supplementary Fig. 5f-h**.

### 10X Genomics Visium Analysis

Raw sequencing data for each tissue section were processed using the SpaceRanger (v1.1.0) pipeline from 10x Genomics. FASTQ files were aligned to the GRCh38 human reference genome (GENCODE v32). To generate the spatial gene expression matrices, the software integrated transcriptomic data with high-resolution .jpeg tissue images of H&E stainings, utilizing slide-specific capture area spot layouts to map gene expression counts to their precise spatial coordinates. This process was applied to all four capture areas per Visium slide, resulting in standardized spot-by-gene count matrices for downstream spatial analysis.

#### Quality Control (QC) and filtering of spots

Downstream analysis of Visium spatial transcriptomics data was conducted using a hybrid workflow leveraging Scanpy v1.9.5 ^(63)^ within Python v3.8.5 and Seurat (v4.0.5). To ensure data integrity, we calculated key quality control metrics for every spatial spot, including total library size (UMI counts), molecular complexity (number of detected genes), and the percentage of reads mapping to mitochondrial and ribosomal genes.

These metrics were visualized spatially to assess samples for signs of over-permeabilization or excessive read diffusion. Following this assessment, low-quality spots were identified and manually removed using the in-house shiny-histopathology application (https://github.com/Single-Cell-Genomics-Group-CNAG-CRG/shiny-pathology). This curation process specifically excluded spots overlapping with histological artifacts, such as tissue folds or regions exhibiting over-exposed (“burned”) staining and necrotic areas delimited by an expert histopathologist by manually annotation in a blinded-manner using QuPath ^(64)^, ensuring that downstream analyses were restricted to high-quality tissue regions.

#### Processing of samples

Following the exclusion of low-quality barcodes, the curated datasets were structured as AnnData objects (v0.9.2) within the Scanpy framework. To correct for technical variations in sequencing depth, gene expression counts were normalized using a total-count scaling approach, adjusting each spatial spot to a fixed library size of 10,000 reads. The data were subsequently log-transformed (log1p) to stabilize variance.

To capture biologically relevant spatial heterogeneity, we utilized SpatialDE v1.1.3 ^(65)^ to identify genes with significant spatial expression patterns. These spatially variable features were scaled and subjected for PCA. A neighborhood graph was constructed using the top 30 principal components, serving as the basis for unsupervised clustering. This initial clustering was performed to validate data coherence, confirming that the resulting transcriptomic clusters spatially recapitulated known histological features and reflected the structural complexity of the tumor tissue.

To enhance the visualization of specific marker genes, we applied the MAGIC v3.0.0 ^(66)^ imputation algorithm. This smoothing approach was used exclusively for spatial visualization in Scanpy to mitigate technical dropout and better resolve regional expression patterns; however, all quantitative analyses were performed using non-imputed data.

### Data analysis

#### RT Signature

The Rhabdoid Tumor (RT) signature was generated comparing bulk RNA-seq data from MRTs samples against other pediatric RNA-seq tumor datasets. Concretely, RNA-seq count matrices were downloaded from the St. Jude Cloud data portal ^(37)^.

Differential expression analysis was performed using the R package DESeq2 (v1.34.0), comparing samples from each ATRT-MRT subtype against samples from all other pediatric tumor types individually, resulting in 37 pairwise comparisons per subtype corresponding to the 37 non-ATRT-MRT tumor types included in the dataset. For each comparison, genes were classified as positively or negatively differentially expressed using predefined thresholds. Positive hits were defined as genes with baseMean >20, log2FC > 1, and p-value < 0.01, whereas negative hits were defined as genes with baseMean > 20, log2FC < ™0.25, and p-value < 0.01.

Broad subtype signatures were defined by aggregating results across all comparisons. Genes were retained if they passed the differential expression filters in at least 30 of 37 comparisons, showed consistent fold-change directionality (positive or negative, respectively) in at least 32 comparisons, and met test statistic consistency thresholds (> ™1.5 for positive hits; < 1.5 for negative hits) in at least 34 comparisons.

The RT signature was used in Multiome and Visium by selecting genes with an absolute mean log2 fold-change (log2FC) > 1.25. To quantify signature activity, we employed a Multivariate Linear Model (MLM) framework, as implemented in decoupleR ^(67)^, using the mean log2FC values as weights for each gene. This scoring approach was applied independently to the single-cell Multiome dataset and the spatial Visium spots.

- Single-Cell Multiome Scoring: Signature scores were calculated (v2.9.1) for the integrated cohort Seurat object. Distribution of these scores across patient-derived Multiome clusters was visualized using ggplot2 (v3.4.4) via violin plots overlaid with jittered data points, stratified by patient ID (**Supplementary Fig. 2a**).
- Spatial Visium scoring: Activity scores were computed (v1.5.0) and mapped onto the tissue coordinates of each sample using Scanpy to visualize the spatial distribution of the RT program (**Supplementary Fig. 2b**).

#### Gene Co-expression Network Analysis

We performed high-dimensional weighted gene co-expression network analysis (hdWGCNA) within the tumor cell population using the R package hdWGCNA v0.4.7 (33) with WGCNA v.1.73 ^(68)^. To ensure more robust results of this network analysis, we first constructed “meta-cells”, small aggregates of transcriptomically similar cells originating from the same tumor, which greatly reduces the inherent sparsity of single-cell transcriptomic data. Furthermore, to perform an unbiased transcriptomic analysis while limiting the contribution of noisy signals, we also filtered out genes that were expressed in fewer than 5 % of cells. We performed a parameter search to assess a suitable soft-power threshold for the network, selecting the smallest soft-power value with a scale-free topology model fit of at least 0.8. We then calculated the weighted gene co-expression network, performed hierarchical clustering on the network, and grouped cells into co-expression modules using the dynamic tree cutting ^(69)^ (R package v1.63.1) algorithm. We calculated module eigengenes (MEs), cell-level gene expression scores, for each of these co-expression modules, and we calculated the eigengene-based connectivity (kME) of each gene by calculating the pearson correlation of gene expression with the overall module expression signature (ME). To visualize the “hub gene” networks, we retained the 250 strongest co-expression links for the top 25 genes ranked by kME per module, and we visualized the resulting network using the R packages tidygraph (v1.3.1) and ggraph (v2.2.2).

Module 1 (M1) is a transcriptional metaprogram characterized by a strong enrichment of oxidative phosphorylation (OXPHOS) genes (**Supplementary Table 3**) and exhibited disproportionately high activity in a subset of patients, particularly those who had undergone prior chemotherapy. Due to its dominant variance, the inclusion of M1 in global scaling obscured the resolution of more subtle transcriptional programs. Given the ambiguity regarding whether this signal represents a specific treatment-induced biological state or a technical artifact, we omitted M1 from the primary visualization in **Fig. 2c** to ensure a balanced representation of the broader metaprogram landscape. However, M1 is fully represented at the pseudobulk level in **Supplementary Fig. 3h**.

#### Regulon Analysis

The same cells that passed the QC and were used for the cohort integration were used in the regulon analysis. Chromatin accessibility data was processed using the pycisTopic (v.2.0a0) package ^(38)^. Pseudobulk profiles were generated based on patient-derived Multiome cluster annotations and open chromatin regions were identified by narrow peak calling using MACS2 (v.2.2.9.1)(70) with the following parameters: --shift 73, --extsize 146, --keep-dup all, and q-value < 0.05. A consensus peak set was subsequently generated using the get_consensus_peaks function with a peak_half_width of 250 bp. To decompose the accessibility matrix, Latent Dirichlet Allocation (LDA) was performed; after evaluating model convergence, a 200-topic model was selected. Topics were binarized using methods ntop (top 3000 regions) and otsu. Differential accessible regions were calculated for patient-derived Multiome cluster annotations.

To explore finer patient-specific differences in gene regulatory networks, a parallel regulon analysis was performed in a patient-specific manner. For this, MACS2 peak calling was executed independently for each patient without establishing a cross-patient consensus peak set. Similarly, LDA model evaluation and topic binarization were optimized for each patient independently based on individual model convergence.

For both cohort and patient-specific workflows, raw transcriptomic data was extracted from integrated Seurat objects and processed using Scanpy (v.1.10.4)^(63)^, including log-normalization. The SCENIC+ (v.1.0a2) pipeline was implemented via Snakemake using the Aerts Lab motif collection (V10 2022: v10nr_clust_public). The pipeline was executed with default parameters, with the exception of biomart_host set to “jan2020.archive.ensembl.org” and a min_target_genes threshold of 20. Extended eRegulons were retained only if no direct eRegulon was identified for a given TF. Dimensionality reduction was performed on the gene-based AUC scores for both direct and extended eRegulons. To evaluate the patient-specific eRegulons specifically regulating HGF or MET, regions from different patients overlapping were unified across patients.

eRegulons selection was based on the overlap with co-expression metaprogram genes. Specifically, eRegulons were selected if at least 20 % of any metaprogram gene set overlapped with the eRegulon target genes.

The cluster specificity of eRegulons across patient-derived Multiome clusters was assessed using the regulon_specificity_scores (RSS) function. Highly specific eRegulons were defined as those above the elbow point of the ranked RSS curve, identified via the KneeLocator function from the kneed library (v.0.8.5)^(71)^.

Gene regulatory networks were visualized adapting the scenicplus.networks functions. To optimize clarity, visualization was restricted to genes linked to the top 100 regions ranked by triplet_rank. Edge weights (line widths) were scaled according to the importance_TF2G score. Regions were classified as Promoter if they were within 1Kb of a TSS, Ensembl enhancers if they overlapped with enhancers in Ensembl regulatory feature annotations (Ensembl regulatory build GRCh38 v115), or >1Kb from Promoter for the rest. For patient-specific HGF and MET networks, edge weights were scaled to the number of patients linked to a specific region.

To visualize the TF regulatory contribution for a specific eRegulon, region-based AUC values for regions linked with the TF were extracted and values were scaled by their importance_x_abs_rho to account for the differential regulatory weights of individual regions within the eRegulon.

#### Deconvolution of Visium spots

To estimate cell-type compositions within the Visium spots, we performed spatial deconvolution using SPOTlight (v2.0.1). We used a patient-matched reference strategy: for each Visium sample, the corresponding single-cell Multiome data from the same patient served as the reference for deconvolution. This approach allowed for the precise mapping of patient-specific cellular states onto their respective spatial coordinates. Concretely, we run SPOTlight to restrict the gene set to the marker genes for each cell population cluster (p-value adj. <0.05 and log2FC>0.25; except p7 which we use log2FC>0.5 to avoid noisy results) along with the corresponding HVG set per sample which further optimizes performance and computational resources.

#### Niche analysis

To characterize the broader tissue architecture, we aggregated the deconvolved proportions of individual patient-derived Multiome clusters into major lineages, including Tumor, Immune, and Other stromal compartments. This consolidation allowed us to map the spatial distribution of primary cell groups across the Visium spots (also see **Fig.1g; Extended Data Fig. 2b**).

To quantify local cellular heterogeneity, we computed Shannon Entropy scores for each Visium spot based on the cell-type proportions obtained from deconvolution. For each spot, we defined a spatial neighborhood including all spots within a 100 µm radius and averaged their cell-type proportions to obtain a local neighborhood composition. Shannon Entropy was then calculated on this neighborhood-level composition, where higher values indicate more heterogeneous, mixed microenvironments and lower values reflect homogeneous, cell-type-dominated regions.

To identify recurrent spatial topological niches, we performed K-means clustering using neighborhood-level features, including local cell-type proportions and Shannon Entropy scores. Using a k-nearest neighbors-based representation and specifying five clusters, this unsupervised approach grouped spatial neighborhoods into distinct niche types reflecting shared compositional and heterogeneity patterns across samples.

#### Cell-Cell Communication

To infer potential intercellular signaling interactions between the identified transcriptomic niches, we used CellChat v2.2.0 ^(72)^. This analysis was performed independently for each patient sample to capture individual-specific signaling between niches.

We employed a spatially constrained approach to prioritize biologically relevant paracrine signaling. Ligand-receptor interactions were inferred using the CellChatDB.human database, with communication probabilities calculated based on the law of mass action. Crucially, to reflect the local nature of signaling within the tissue, we restricted the analysis to a maximum interaction distance of 250 µm (approximately 2-3 spots from the source). Signaling pathways were considered significant if the permutation-based p-value was < 0.05.

For heatmap visualization in **Fig. 4e**, for each patient, communication probabilities (interaction strength metric, not a literal probability) were normalized to the maximum observed value to allow for inter-sample comparison.

#### Analysis of CRISPR-Cas9 from DepMap

TPM expression and gene effect (Chronos) of *MET* were obtained from the DepMap Portal (DepMap Public 26Q1; https://depmap.org/portal). Cell lines were filtered by Pediatric model type and those ones without a gene effect score were excluded. Based on the Oncotree Primary disease, the 224 cell lines were stratified into ‘ATRT’, ‘RTK’, ‘ERRT’ and ‘other childhood’ cell lines. For scatter plot visualization in **Supplementary. 7c**, cell lines were grouped into low (<5 log2(TPM+1)) and high (> 5 log2(TPM+1)) MET expressing groups.

#### Module Scoring in St Jude Cloud Cohort

Feature count matrices and companion metadata from bulk RNA-seq data were downloaded from St. Jude Cloud Genomics Platform for all available tumor samples as of the 13th of April 2026^(37)^.

Raw count matrices were imported and merged into a unified AnnData object (v0.9.2), with sample-specific clinical annotations appended as metadata. The dataset was filtered to restrict analysis strictly to primary samples collected at diagnosis; all relapsed, metastatic, and patient-derived xenograft (PDX) samples were excluded. To facilitate comparative analysis with malignant rhabdoid tumors (MRTs), the detailed diagnostic classifications provided by the St. Jude Cloud were manually adjusted into intuitive, standardized acronyms representing major disease entities.

A representative subset of pediatric malignancies was selected for downstream comparison, spanning hematological malignancies (Acute Myeloid Leukemia [AML], B-cell Acute Lymphoblastic Leukemia [B-ALL], and T-cell Acute Lymphoblastic Leukemia [T-ALL]), major sarcoma groups (Ewing Sarcoma [EWS], Osteosarcoma [OS], and Rhabdomyosarcoma [RMS]), and neural-derived embryonal tumors (Diffuse Midline Glioma [DMG], Neuroblastoma [NB], and Medulloblastoma [MB]). This filtration process yielded a final cohort of 3029 samples. Raw expression counts were subsequently normalized to logCPM.

To quantify transcriptional module activity across the bulk St. Jude cohort, we used reference weights derived from our single-cell dataset. Specifically, logCPM pseudobulk expression profiles were generated from the single-cell tumor fraction, and the average gene expression across all patients was calculated for the genes comprising each target module. These average expression values were then used as feature weights for Multivariate Linear Model (MLM) analysis, implemented via the decoupleR package (v1.5.0), to compute robust module activity scores for each sample in the St. Jude dataset.

### Establishment of patient-derived MRT models

A small fraction of fresh or cryopreserved tumor fragments were used to establish patient-derived MRT cell lines from p9, p10, p11, p13 and p19 biopsies (see ‘*Patient specimens and biobanking*’ section). For p5 and p15 cell lines, an intermediate in vivo amplification step was required prior to in vitro cell line derivation. Following the harvest of xenograft tumors, samples were processed for in vitro culture. Briefly, tumor samples were minced into small fragments (< 0.5 mm^3^) using sterile scalpels in pre-warmed RPMI 1640 medium. To dissociate the tissue, fragments were enzymatically digested with 200 U/mL collagenase (Sigma, C5138), 160 U/mL DNase I (Sigma, D4263) in RPMI 1640 for 30 min at 37°C under gentle agitation. Following a washing step, the digested mixture was further dissociated using 0.25 % Trypsin-EDTA (Gibco, 25200056) at 37 °C for 5 min. After trypsin inactivation, the mixture was filtered through a 100-μm cell strainer and equal aliquots were cultured in parallel in both tumor Stem Media (TSM) base ^(73)^ and in TSMserum medium (TSM base substituting B27 supplement with 10 % FBS). Upon confirmation of *SMARCB1* genomic alterations and validation of protein loss via Western blot (**Fig. 5d**), mycoplasma-free cells exhibiting optimal growth were expanded, cryopreserved in Synth-a-Freeze cryopreservation medium and stored in liquid nitrogen for future use.

### Cell culture

CHLA-02-ATRT (RRID:CVCL_B045; male), CHLA-05-ATRT (RRID:CVCL_AQ41; male), HFF-1 (RRID:CVCL_3285; male), SH-SY5Y (RRID:CVCL_0019; female) and HEK293 (RRID:CVCL_0045; female) cells were purchased from the American Type Culture Collection (ATCC) and maintained according to the repository’s instructions. Patient-derived rhabdoid tumor cells were cultured in TSMserum medium, except for p13 (ATRT-MYC) cells that were cultured in DMEM/F12 (Gibco, 21331020) supplemented with 1 % B27 supplement (Gibco, 17504044), 20 ng/mL Human EGF Recombinant protein (Gibco, PHG0311), 20 ng/mL Human FGF-basic Recombinant Protein (Gibco, PHG0026) and 100 U/mL penicillin-streptomycin (Gibco, 15140122). Fresh growth media was added every 48 to 72 hours, and cultures were maintained at 37°C in a humidified incubator at 5 % CO_2_. Absence of mycoplasma contamination was routinely tested using the EZ-PCR Mycoplasma Detection kit (Biological Industries, 20-700-20) according to the manufacturer’s instructions. To minimize phenotypic drift, all experiments were conducted using cell lines below passage 20.

### Quantitative Real-Time PCR

Cells were harvested in TRIzol (Thermo, 15596018) and RNA was purified using the RNeasy Mini kit (QIAgen, 74106) following the manufacturer’s instructions. 1 µg of total RNA was reversely transcribed to cDNA with the High-Capacity cDNA Reverse Transcription Kit (Applied Biosystems, 4368814) according to the manufacturer’s instructions. Human TaqMan probes for *HGF* (Hs.PT.58.81779) and *MET* (Hs.PT.58.339430) were purchased from Integrated DNA technologies, and relative expression levels were calculated using the 2^™ΔΔCT^ method ^(74)^ and normalized to *GAPDH* (Hs.PT.39a.22214836) expression levels.

### Western blotting

For protein extraction, cells were scrapped and lysed in RIPA Lysis Buffer (Thermo, 89901) containing 1x EDTA-free Protease Inhibitor Cocktail (Roche, 11873580001), 1x phosphatase Inhibitor Cocktail 2 and 3 (Sigma, P5726 and P0044, respectively), and protein was quantified using the Pierce™ BCA® Protein Assay Kit (Thermo, 23225). 20 µg protein were separated on 4-12 % Bis-Tris gels (Thermo, NP0323BOX), transferred to pre-activated PVDF membranes (Cytiva, 10600029) and blocked in 3 % BSA in TBS-T (blocking solution) for 1 hour. Membranes were incubated at 4°C overnight in α-SMARCB1 (Abcam, ab222519; 1:1,000), α-GAPDH (CST, 2118; 1:1,000), α-MET (CST, 8198; 1:1,000), α-phospho-MET (Tyr1234/1235) (CST, 3077; 1:1,000) primary antibodies diluted in blocking solution, washed thrice in TBS-T, and incubated with α-rabbit-HRP-linked antibody (1:5,000; Dako, P0399) for 1 hour at room temperature. After three washes in TBS-T, membranes were developed with SuperSignal™ West Pico PLUS Chemiluminescent Substrate (Thermo, 34577) in an iBright CL1500 Imaging system (Invitrogen, USA). For reprobing, membranes were stripped of previously bound antibodies by incubating in stripping buffer (Thermo, 46430) for 20 min following the manufacturer’s instructions. Membranes were blocked and reprobed as previously described.

### IC_50_ determination

To determine the IC_50_, a dose-response assay was performed using DMSO or 5-fold serial dilution of PHA-665752 (MedChem Express, HY-11107), ranging from 1.28 nM to 100 µM. Briefly, cells were seeded (2-4 × 10^3^ cells/well) in a white-wall, clear bottom 96-well plate (SPL Life Sciences, 33696) and allowed to adhere overnight. The day after, cells were treated with the corresponding 2x treatment condition. Cell viability was evaluated using CellTiter-Glo Luminescent Cell Viability Assay (Promega, G7571). Cells were lysed using 100 µL of reconstituted CellTiter-Glo directly on the plate, and incubated for 20 min in an orbital shaker (100 RPM, RT). The luminescence signal was measured in a Synergy H1M multi-mode microplate reader (BioTek Instruments, USA). For data analyses, background luminescence was subtracted from all experimental wells. Data were normalized to the vehicle control (DMSO) and IC_50_ values were determined using non-linear regression analysis of the dose-response curves in GraphPad Prism software v10.4.1 (GraphPad Software, USA).

### Annexin V/DAPI analyses

To assess the cytotoxic effects of the PHA-665752 MET inhibitor, HFF-I, p5 and p13 cell lines were used. Cells were treated with 50 nM, 150 nM, 300 nM and 900 nM PHA-665752, or the vehicle at maximum concentration. After 24 hours of incubation, cells were harvested and stained with Annexin V-Alexa Fluor® 647 (1:25; Biolegend, 640912) and 0.5 μg/mL 4′,6-diamidino-2-phenylindole (DAPI) (Thermo, 62248) in Annexin Binding buffer (10 mM HEPES pH 7.4, 150 mM NaCl (Sigma, S5886), 2.5 mM CaCl_2_ (Sigma, C3306) in PBS pH 7.4) and kept on ice. Sample acquisition was performed within one hour of preparation using a CytoFlex SRT flow cytometer (Brand, Country), and flow cytometry data was analyzed using FlowJo software v10.10.0 (BD Biosciences, USA) to quantify the percentage of necrotic and early-late apoptotic cells.

### Neutralizing antibody assay

To evaluate the therapeutic potential of ficlatuzumab (MedChem Express, HY-P99196), cells were seeded as previously described and allowed to adhere overnight before treatment with 4-fold serial dilutions, ranging from 1,000 μg/mL to 244 ng/mL. Following a 96 hours incubation, cell viability was measured and analyzed as previously described (see ‘*IC*_*50*_ *determination*’ section). To account for non-specific antibody effects, raw luminescence signal was background-substracted and subsequently normalized to Human IgG1 (MedChem Express, HY-P99995) Isotype control at 1,000 μg/mL.

### Drug perturbation studies

#### Compound preparation

Compounds from the IRB compound library were stored as 10 mM stock solutions in DMSO and pre-dispensed into 384-well destination plates (384-well culture plate, Revvity) using an Echo® 650 acoustic dispenser (Labcyte, Beckman Coulter). Acoustic dispensing volumes were calculated to achieve the desired final compound concentrations in the assay. In all experiments, a constant final concentration of vehicle was maintained across all wells and never exceeded 0.5 % to avoid solvent-related toxicity. After compound pre-printing, plates were sealed and stored at ™20 °C until use.

#### Cell viability assay and compound screening

Cells were harvested by standard trypsinization for adherent cells or by Accutase for cells growing in suspension, counted, and diluted in the appropriate culture medium for each cell line to reach the desired final seeding density (**Supplementary Table 5**). Compound-containing plates were equilibrated at room temperature for 30 min prior to seeding, and 50 µL of cell suspension were added per well using an automated Multidrop™ dispenser (Thermo Fisher Scientific). Plates were then incubated for 72 h at 37 °C and 5 % CO_2_.

Cell viability was assessed using the CellTiter-Glo® Luminescent Assay. After incubation with compounds, plates were equilibrated at room temperature for 30 min and 15 µL of CellTiter-Glo reagent were added per well using a Multidrop™ system integrated into the automated Cell Explorer platform (Revvity, USA). After 10 min of incubation, luminescence was measured using an EnVision® Xciteplate reader (Revvity, USA). The luminescent signal is proportional to intracellular ATP levels and therefore to the number of metabolically active cells.

Optimal seeding density for each cell line was determined in preliminary dose–response experiments using three cytotoxic compounds at different cell densities (data not shown). Conditions providing an adequate dynamic range and reproducible responses were selected for subsequent experiments.

Once optimal seeding densities were established, a total of 44 compounds were selected from the IRB focused libraries acquired from MedChemExpress based on data from our analyses and literature evidence. These included 21 FDA-approved compounds, 12 kinase inhibitors, 2 compounds from the immunology/inflammation subset, 4 compounds in clinical trials, 1 natural product, and 1 protein–protein interaction modulator. Compounds were tested in all 12 cell lines at final concentrations of 1,000 and 100 nM, in triplicate, following the viability assay protocol described above.

Assay quality was assessed through the inclusion of internal controls and a randomized experimental design. Compounds and their triplicates were randomly distributed across plates to minimize positional effects. Each plate contained 32 no-drug wells with DMSO as negative controls and 32 maximum inhibition wells containing PHA-665752 at a final concentration of 10 µM, enabling data normalization and calculation of percentage inhibition. All plates yielded Z′ factor values above 0.5, indicating a robust assay with adequate signal separation between positive and negative controls. The coefficient of variation (%CV) of the high-signal control wells (DMSO-treated wells) was below 10 % across all plates, reflecting low intra-plate variability and high overall screening quality. Data analysis was performed using GraphPad Prism, Studies, and Vortex (Dotmatics).

### Animal maintenance

All animal procedures were performed at the Animal Facility of the Parc Científic de Barcelona (PCB). Experimental protocols were approved by the Animal Experimentation Ethics Committee of the PCB (Comitè Ètic d‘Experimentació Animal, CEEA-PCB-24-042). Procedures were conducted in accordance with Directive 2010/63/EU, Spanish Royal Decree 53/2013 and under all applicable institutional guidelines.

Mice were maintained under specific pathogen-free conditions in individually ventilated cages, with a 12h light /12h dark cycle, controlled temperature and relative humidity, and ad libitum access to food and water. Animal health was monitored at least three times per week. A humane endpoint was defined when tumor size exceeded 1500 cm^3^.

### Patient-derived xenograft assays

For patient-derived xenograft (PDX) assays, the indicated cell models were cultured as specified above, washed in DPBS and resuspended in ice-cold DPBS mixed 1:1 with growth-factor reduced matrigel (Cultek, 346231). 20-week-old NOD.Cg-*Prkdcscid Il2rgtm1WjI*/SzJ NSG mice (Envigo; balanced 1:1 male to female ratio) were inoculated subcutaneously in the right flank with 1 × 10^6^ cells in 100 μL total volume. Tumor dimensions were measured three times per week from day 8 post-inoculation using digital calipers. Tumor volume was calculated as Volume = (Length × Width^2^)/2, where length represents the longest tumor dimension and width the perpendicular dimension. On day 14 post-inoculation (mean tumor volume: 120 mm^3^), mice were randomized into vehicle or treatment groups, balanced by sex distribution.

### Crizotinib formulation and administration

Crizotinib (MedChem Express, HY-50878) was received in powder. To enhance drug solubility, 1 % DMSO was added to create an homogeneous mixture, and the remaining volume was gradually added using a common vehicle consisting of 0.5 % Carboxymethylcellulose sodium salt (Merck, C5013) in 0.9 % NaCl (S9625, Merck) in water. Crizotinib and the corresponding vehicle control were freshly prepared daily. Mice received 75 mg kg^-1^ or equivalent volume of vehicle once daily by oral gavage at a dosing volume of 10 mL kg^-1^.

### Declaration of generative AI and AI-assisted technologies in the manuscript preparation process

During the preparation of this work, the authors used ChatGPT (OpenAI, GPT-5.3) and Gemini 3 Flash (Google) in order to assist with code refinement, and language editing to improve readability. After using these tools, the author reviewed and edited the content as needed and take full responsibility for the content of the published article.

