## Supplementary Data Figures for "MET inhibition exploits conserved regulatory dependencies in malignant rhabdoid tumors"

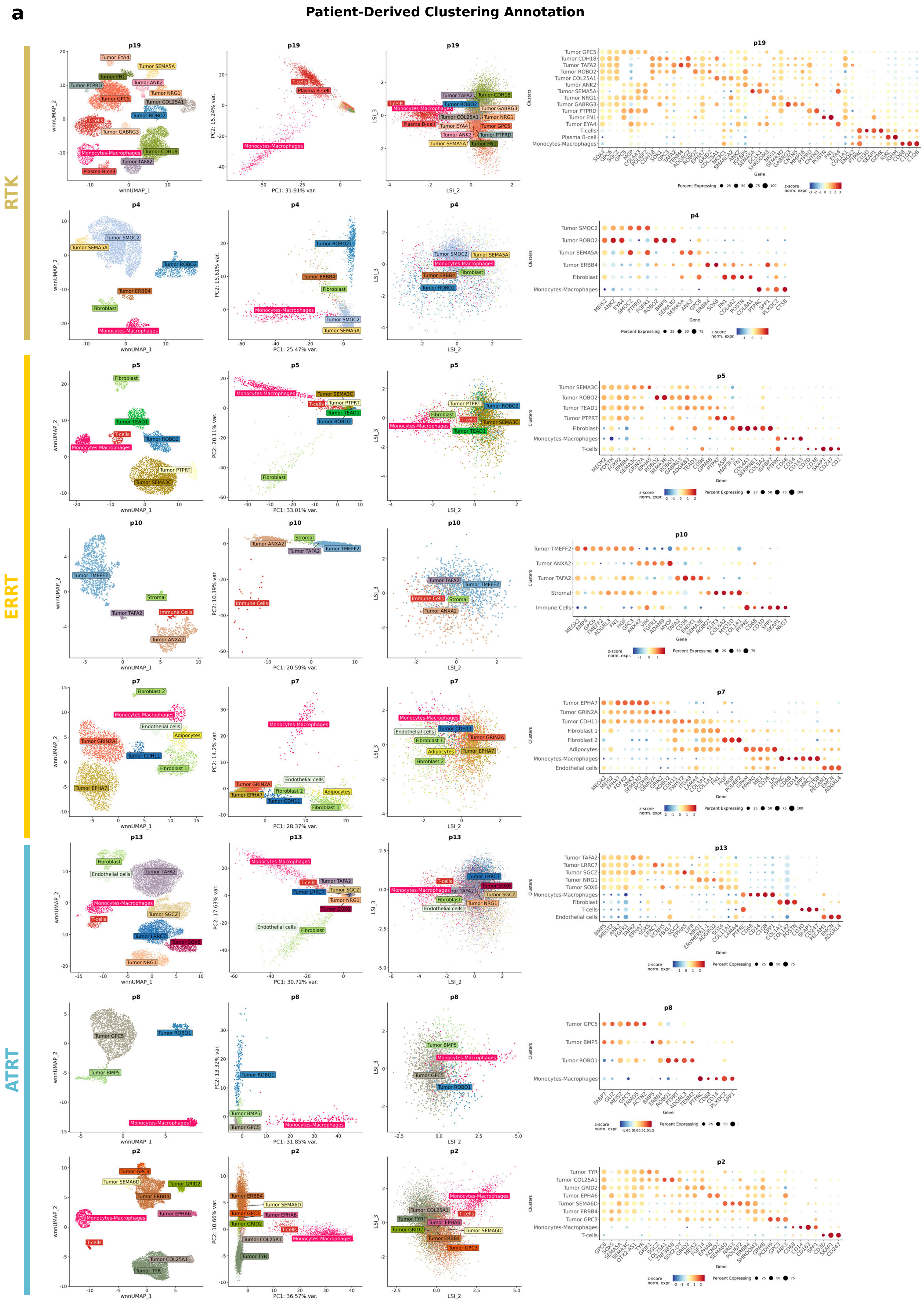


Supplementary Fig. 1

**Supplementary Fig. 1 | Annotation of patient-derived Multiome clusters.**

**(a)** From left to right: UMAP representing the annotated cell clusters; RNA-based scatter plot with PC1 (x-axis) and PC2 (y-axis) cells colored by the annotated clusters; ATAC-based scatter plot with LSI2 (x-axis) component and LS3 component (y-axis) cells colored by the annotated clusters; Dot plot illustrating the expression of top marker genes used to define and annotate each cell cluster. To enhance signal stability, raw counts were aggregated into pseudobulks per cluster, normalized to log-CPMs, and z-scored across all groups. The color intensity represents the relative expression level (z-score), while the dot size denotes the percentage of cells within each cluster expressing the gene.

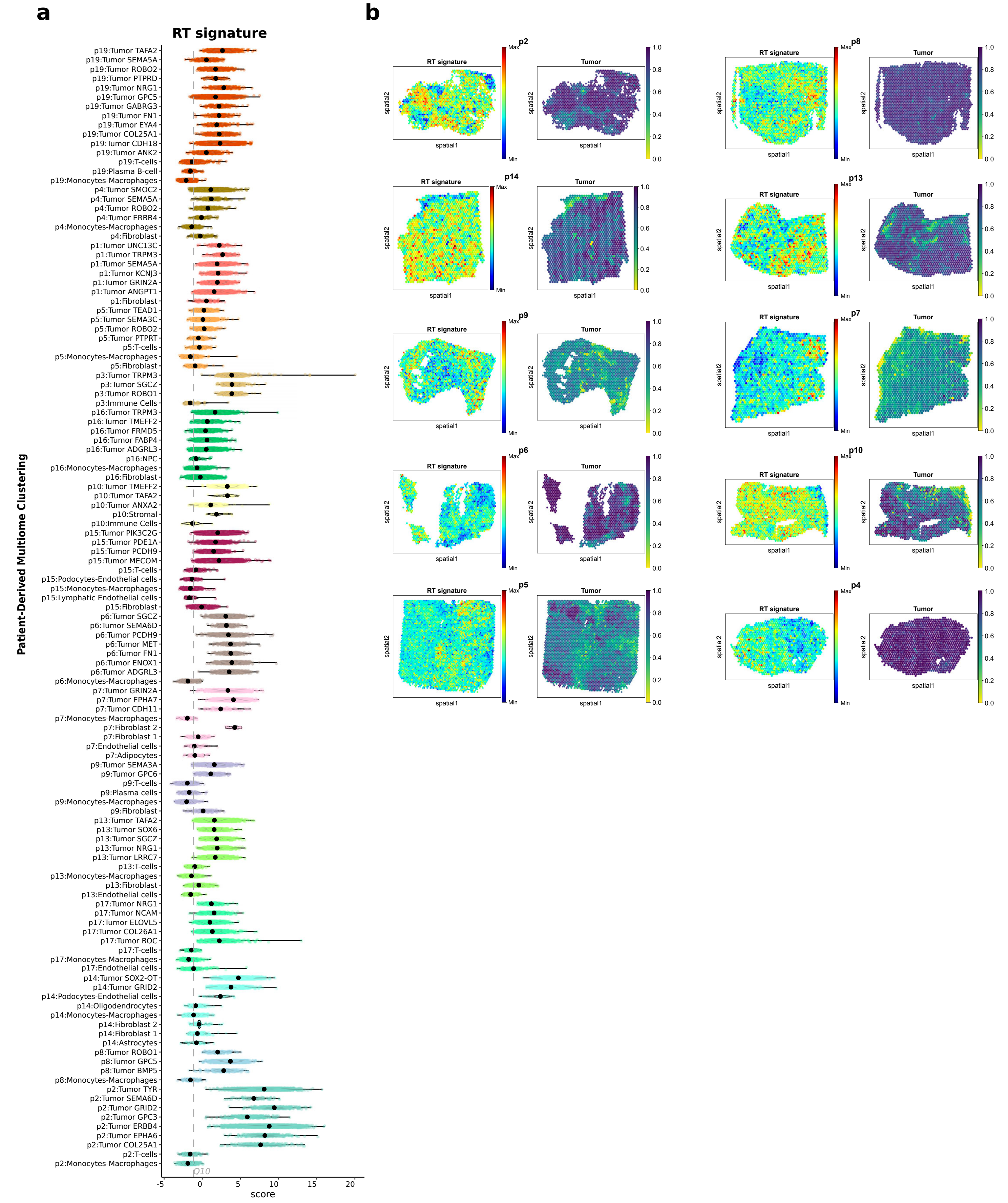


Supplementary Fig. 2

**Supplementary Fig. 2 | Annotation of malignant rhabdoid cells. (a)** Violin plot representing the distribution of the rhabdoid tumor signature scores in single-cell data across patient-derived Multiome clusters. Violin plots and overlapping data points are colored by patient identity. Dashed line represents the 10th percentile of the distribution of all patient cell scores. **(b)** Spatial distribution of the RT signature within tissue sections. Left: Spatial maps of Visium spots colored by RT signature activity, scaled to the relative minimum and maximum values per sample. Right: Spatial deconvolution results showing the estimated proportion of malignant tumor cells per spot.

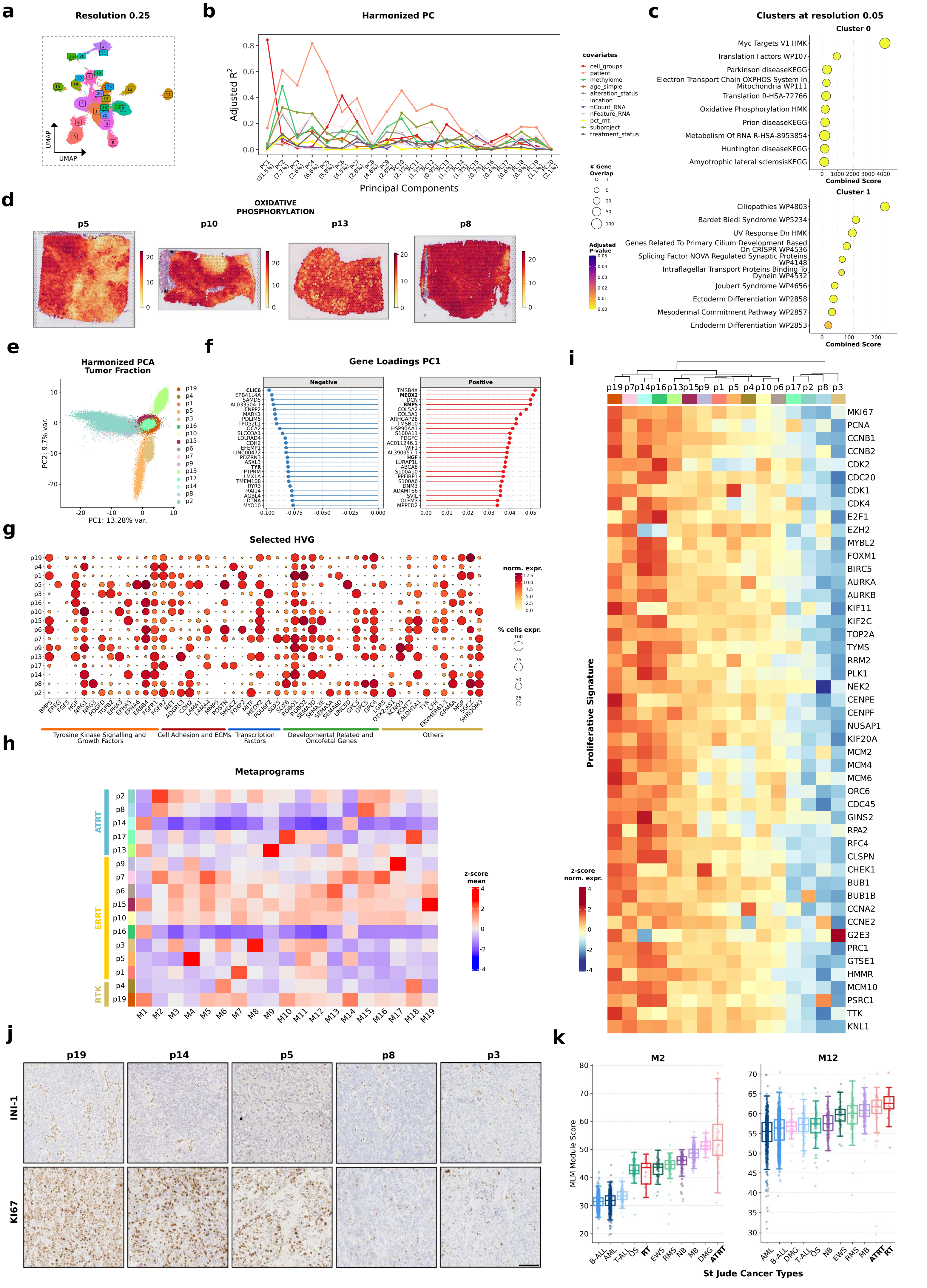


Supplementary Fig. 3

**Supplementary Fig. 3 | Inter-patient variability and co-expression metaprograms across rhabdoid tumors. (a)** UMAP of integrated rhabdoid patient samples color-coded by unsupervised clustering at increased 0.25 resolution. **(b)** Principal component regression of the first 20 PCs corrected by Harmony and metadata covariates. Y-axis represents the adjusted r2. **(c)** Over-representation analysis of markers for Cluster 0 and Cluster 1 at resolution 0.05; biological terms are ranked by combined score (x-axis), with dot size representing gene count and color indicating the adjusted p-value. **(d)** Spatial distribution of the OXPHOS signature activity (Hallmarks, MSigDB) across spatial samples for patients p5, p10, p13, p8. **(e)** PCA plot of the tumor compartment following Harmony-based integration; cells colored by patient ID. **(f)** Top gene loadings contributing to PC1 variance. **(g)** Dot plot illustrating the pseudobulk expression profiles of selected HVGs from the tumor fraction, categorized by biological function; color intensity represents log-CPM and dot size denotes the percentage of cells expressing each gene (HVG, highly variable genes). **(h)** Heatmap of 19 co-expression-derived metaprograms within tumor populations; columns represent modules and rows represent patients, with scores calculated as the z-scaled mean activity per patient. **(i)** Heatmap showing the relative expression of proliferation-associated genes across patients (rows), ordered by hierarchical clustering. **(j)** Representative INI-1 and Ki67 IHC images of primary tumor biopsies. Scale bar, 100 µm. **(k)** Box plot with the MLM signature score of the gene modules in a selection of cancer types from St. Jude Cloud bulk RNA-seq cohort (n=3029). AML (Acute Myeloid Leukemia), B-ALL (B-cell Acute Lymphoblastic Leukemia), T-ALL (T-cell Acute Lymphoblastic Leukemia), EWS (Ewing Sarcoma), OS (Osteosarcoma), RMS (Rhabdomyosarcoma), DMG (Diffuse Midline Glioma), NB (Neuroblastoma), MB (Medulloblastoma), and RT (Extracranial Rhabdoid Tumor), ATRT (Atypical Teratoid Rhabdoid Tumor).

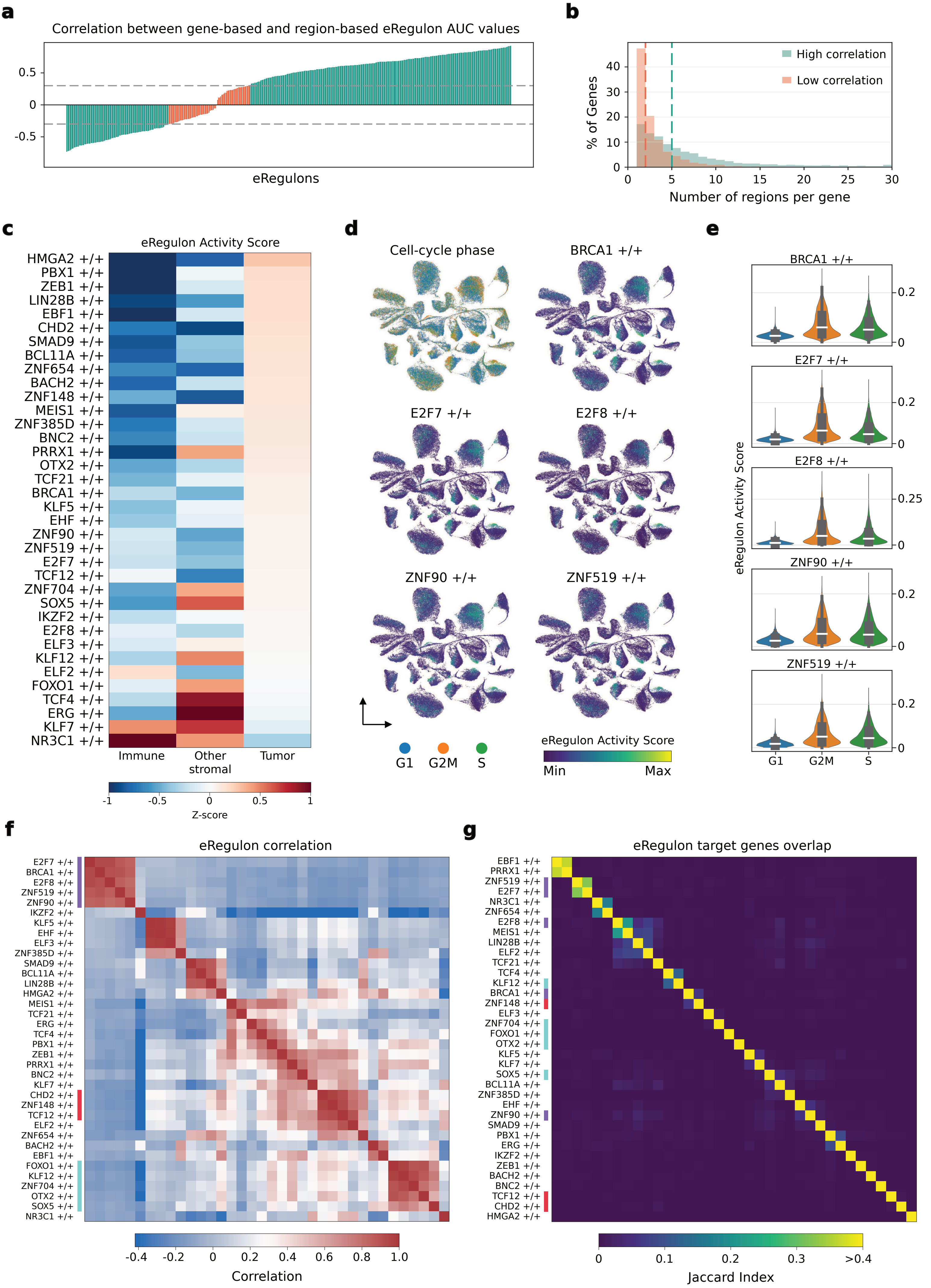


**Supplementary Fig. 4**

**Supplementary Fig. 4 | SCENIC+ eRegulon analysis. (a)** All SCENIC+ called eRegulons ranked by their gene-based and region-based AUC correlation. eRegulons above 0.3 or below -0.3 are considered highly correlated. **(b)** Distribution of the number of regions linked to genes in high and low correlation eRegulons. **(c)** Gene-based eRegulon activity score aggregated for the three major cell groups for the eRegulons selected in Fig. 3a. Z-score was calculated for each cell and aggregated by cell-groups. **(d)** UMAP dimensionality reduction based on gene-based AUC eRegulon scores showing the inferred cell-cycle phase and the eRegulon score of M14 associated eRegulons. **(e)** Violin plot quantification of eRegulon activity enrichment of M14 associated eRegulons shown in **d** in G1, G2M and S cell cycle phases. **(f)** Correlation of gene-based eRegulon scores between selected eRegulons. **(g)** Jaccard index overlap score between selected eRegulons target genes. eRegulons highlighted in **Fig. 3a**,**b** are also highlighted in **f** and **g**.

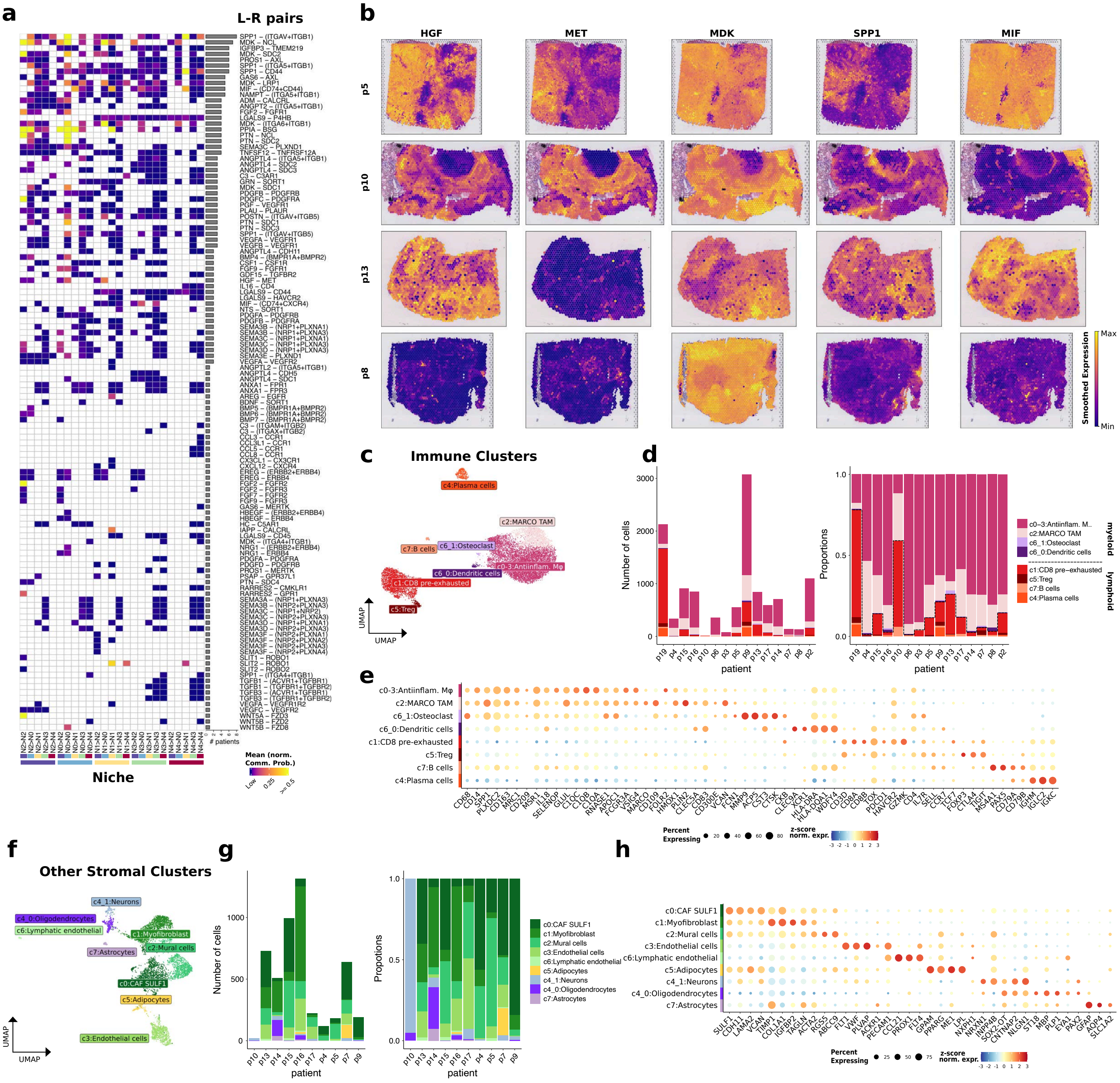


Supplementary Fig. 5

**Supplementary Fig. 5 | Signaling interactions and TME. (a)** Heatmap illustrates the average communication probability between specific source and target tissue niches across the patient cohort. For each patient, raw signaling probabilities (derived from CellChat) were normalized to the maximum observed value to allow for inter-sample comparison. The heatmap displays the mean of these normalized probabilities for each ligand-receptor pair across all patients (ligands can have multiple receptors and vice versa). To emphasize biological signals, the color scale is capped at 0.5 (maximum intensity). Bar plot indicates the total number of patients in whom significant communication was detected for that specific ligand-receptor pair. **(b)** Spatial maps of Visium spots showing the smoothed expression of HGF/MET axis and other specific secreted ligands (MDK, SPP1 and MIF) of interest. **(c)** UMAP showing the integration of immune cells colored by the annotated cell types after unsupervised clustering. **(d)** Left: Bar plot showing the total number of immune cells per patient colored by annotated cell types. Right: stacked barplot with the proportion of immune cell types per patient. **(e)** Dot plot illustrating the expression of top marker genes used to define and annotate each immune cell cluster. To enhance signal stability, raw counts were aggregated into pseudobulks per cluster, normalized to log-CPMs, and z-scored across all groups. The color intensity represents the relative expression level (z-score), while the dot size denotes the percentage of cells within each cluster expressing the gene. **(f)** UMAP showing the integration of other stromal cells colored by the annotated cell types after unsupervised clustering. **(g)** Left: Bar plot showing the total number of other stromal cells per patient colored by annotated cell types. Right: stacked barplot with the proportion of other stromal cell types per patient. **(h)** Dot plot illustrating the expression of top marker genes used to define and annotate each stromal cell cluster. To enhance signal stability, raw counts were aggregated into pseudobulks per cluster, normalized to log-CPMs, and z-scored across all groups. The color intensity represents the relative expression level (z-score), while the dot size denotes the percentage of cells within each cluster expressing the gene.

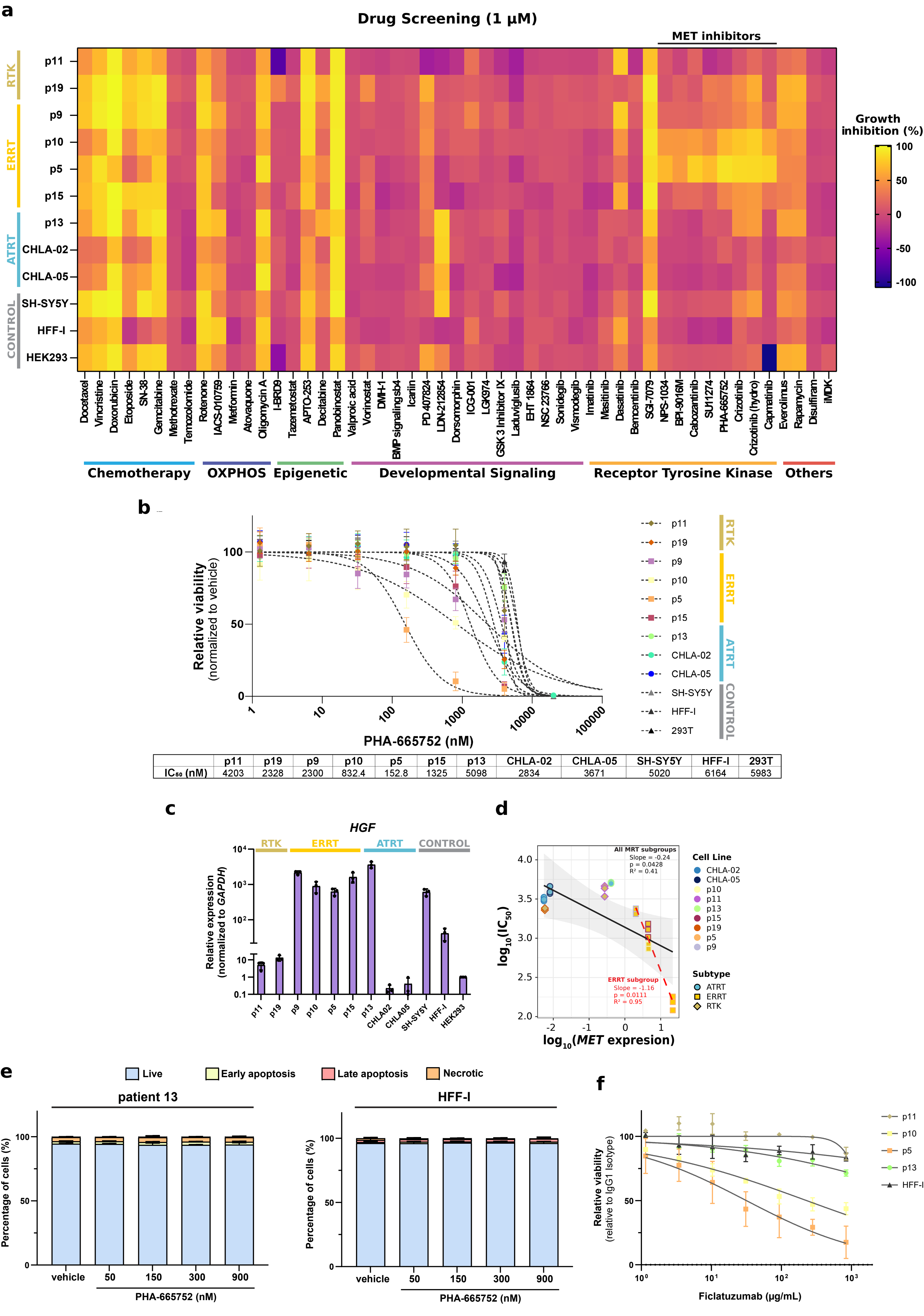


| | p11 | p19 | p9 | p10 | p5 | p15 | p13 | CHLA-02 | CHLA-05 | SH-SY5Y | HFF-I | 293T |
|---|---|---|---|---|---|---|---|---|---|---|---|---|
| $IC_{50}$ (nM) | 4203 | 2328 | 2300 | 832.4 | 152.8 | 1325 | 5098 | 2834 | 3671 | 5020 | 6164 | 5983 |

Supplementary Fig. 6

**Supplementary Fig. 6 | Drug screen at high concentration and validation of MET inhibition. (a)** Heatmap depicting the percentage of growth inhibition across a panel of patient-derived (RTK, ERRT, ATRT) and control (SH-SY5Y, HFF-1 and HEK293) cell lines. Straight line indicates inhibitors targeting MET. Data represented is the mean of three technical replicates following 72 hours of treatment at 1 μM of the indicated compounds (categorized by their primary mechanism of action). **(b)** Cytotoxicity curves of cells treated with PHA-665752 MET inhibitor showing viability relative to the vehicle at 72 hours. Mean $IC_{50}$ values per cell line are indicated (error bars represent SD, n = 3 independent experimental replicates). **(c)** Relative mRNA expression of *HGF* in tumor cell lines assessed by RT-qPCR (n = 3 independent experimental replicates). **(d)** Scatter plot of $log_{10}$ *MET expression* versus $log_{10}$ $(IC_{50})$ values for the PHA-665752 inhibitor. A mixed-effects model was used to evaluate the relationship across all rhabdoid cell lines (black line), and specifically within the ERRT group (red dashed line). The shaded area represents a 95% confidence interval. **(e)** Quantification of live, necrotic, early and late apoptotic patient 13 (left) and HFF-I (right) cells after treatment with vehicle, 50 nM, 150 nM, 300 nM or 900 nM of PHA-665752 inhibitor for 24 hours (errors bars represent SD, n = 3 independent experimental replicates). **(f)** Relative viability of the indicated cell models treated with Ficlatuzumab neutralizing antibody (anti-HGF) for 96 hours, normalized to isotype control (error bars represent SD, n = 3 independent experimental replicates).

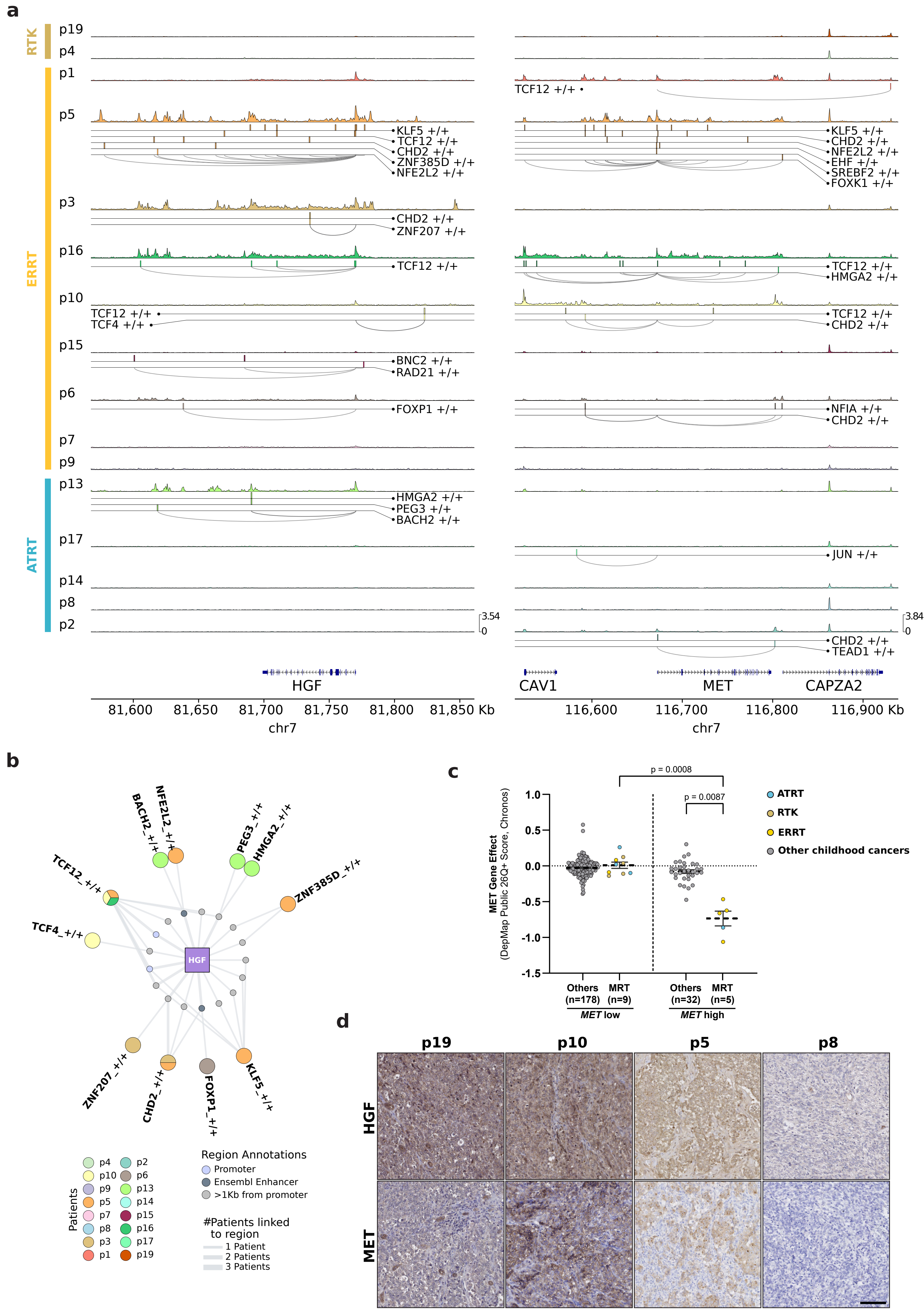


Supplementary Fig. 7

**Supplementary Fig. 7 | Epigenetic and regulatory landscape of the HGF/MET axis and its dependency in rhabdoid tumors. (a)** Patient-specific eRegulon landscape associated with HGF or MET. Tumor fraction pseudobulk scATAC tracks with normalized accessibility shown for all patients. Genomic regions associated with the regulation of HGF or MET are represented as boxes and linked to the promoter with an arch. **(b)** Regulatory network showing the patient-specific eRegulons linked to HGF. Each eRegulon is depicted as a pie-plot showing in which the patients that eRegulon is linked to HGF. Dots representing regulatory regions are classified as *Promoter* if they are within 1Kb of the TSS, *Ensembl enhancers* if they overlap with enhancers in Ensembl regulatory feature annotations, or *>1Kb from Promoter* for all others. Connecting lines width represents the number of patients associated with a specific region. **(c)** Scatter plot comparing dependency scores for MET in childhood cancer cell lines (dots; n = 224). Studied groups were divided into high (> 5 $\log_2$(TPM+1)) and low (< 5 $\log_2$(TPM+1)) *MET* expression, and further annotated by malignant rhabdoid tumor subgroup: ATRT (blue), RTK (brown), ERRT (yellow). Statistical analysis was performed using a One-Way ANOVA with Dunnett's Post-Hoc test. Data are presented as mean ± SEM. **(d)** Immunohistochemistry for HGF and MET on primary tumor samples for p19, p10, p5 and p8. Scale bar, 100 µm.
